# Computational Design of Miniprotein Binders Targeting the Viral Entry in Chikungunya and Dengue, Neglected Tropical Diseases

**DOI:** 10.1101/2025.10.06.680738

**Authors:** Danilo Kiyoshi Matsubara, Peter Park, Iolanda Midea Cuccovia, Hernan Chaimovich

## Abstract

Neglected Tropical Diseases (NTDs) continue to be a global health concern, affecting millions worldwide. Arboviruses such as Chikungunya virus (CHIKV) and Dengue virus (DENV) are particularly threatening due to the lack of specific antiviral treatments, the continual emergence of new variants, and their mosquito vectors. These challenges underscore the urgent need for novel antiviral strategies. Here, we apply computational methods to design miniprotein binders targeting critical viral entry components: the CHIKV envelope, and DENV fusion loop. Our protocol integrates classical and deep learning-based approaches and can be generalizable to other protein targets. Promising binder candidates were further evaluated through all-atom Molecular Dynamics simulations to assess structural stability and binding interactions. We provide complete transparency and a detailed rationale for each design step to facilitate adaptation of these protocols to other neglected disease targets. While the miniproteins presented here represent promising antiviral leads, experimental validation remains essential to confirm their biological activity.

## 1. Introduction

Neglected tropical diseases (NTDs) remain a major global health challenge, imposing substantial morbidity and mortality worldwide [1], [2], especially in low- and middle-income countries [3], [4]. Despite advances in vaccines and antiviral therapies, many viral pathogens still lack effective treatments, and the emergence of new variants reinforces the urgent need for innovative therapeutic strategies [5].

Among viral NTDs, arboviruses such as Chikungunya virus (CHIKV) and Dengue virus (DENV) are of particular concern in tropical and subtropical regions [6], [7]. CHIKV is associated with debilitating joint pain that may persist for years [8], whereas DENV infects millions worldwide annually, with risks of severe dengue and mortality [9]. Yet, no specific therapeutics for treatment exist for either virus [10], [11].

Current antiviral discovery efforts have focused on small molecules [12], [13], [14] and monoclonal antibodies [15], [16]. Small-molecule inhibitors often target non-structural proteins, including viral polymerases and proteases [17], [18], which contain defined active sites with distinct binding pockets, making them ideal for inhibition by small-molecule compounds [19].

Another strategy is to block viral entry by targeting structural proteins. For instance, CHIKV uses its envelope (E1-E2) to bind one of its entry receptors, MXRA8 [20], facilitating attachment and initiating infection. In the case of DENV, although multiple receptors can mediate viral entry, the process converges in the endosome, where its fusion loop, exposed by the low pH, mediates membrane fusion with the host cell [21]. Therefore, the CHIKV envelope/MXRA8 interface, and the DENV fusion loop/endosome membrane interaction represent attractive entry-inhibition targets [22], [23].

However, such interaction interfaces are often large, relatively flat, and lack well-defined pockets, making them difficult to inhibit with small molecules [24]. Even when the surfaces are small and convex, as in the DENV fusion loop, the absence of well-defined binding pockets presents similar challenges. Protein-based therapeutics can offer higher specificity and broader interaction surfaces, making them more suitable for disrupting these types of molecular interactions [25].

Monoclonal antibodies have shown promising results, but concerns remain regarding antibody-mediated enhancement (ADE) in DENV [26], along with issues of high costs [27], [28], which can limit accessibility, stability [29] and reproducibility [30], [31]. De novo designed miniprotein binders provide an attractive alternative [32], [33]. Unlike natural proteins, these molecules are designed from scratch [34], allowing customization of size, topology, and binding interfaces. They can also be expressed in bacteria at scale, offering cost and production advantages.

Computational protein design has already demonstrated promising results in generating binders against viruses such as SARS-CoV-2 [35], relying on energy-based approaches to sample sequences and conformational space. Recent advances in deep learning-based structure prediction and protein design [36], [37] have further accelerated the development of more efficient and accurate design pipelines.

Building on these advancements, this work presents a general protein design protocol for generating miniprotein binders targeting key viral entry proteins of CHIKV (envelope), and DENV (envelope - fusion Loop). The aim is to develop miniproteins with potential as antiviral therapeutics or as starting points for future optimization and refinement. To evaluate the designed binders, Molecular Dynamics (MD) simulations were performed on selected candidates, both in complex with their targets (to assess interaction stability), and in solution (to evaluate the structural stability). The rationale behind each protocol step and the challenges encountered are also discussed, providing a reference for researchers interested in adopting or adapting similar design strategies.

## 2. Methods

The protocol consists of two distinct phases (Fig. 1). The first, referred to as *Model Generation* (Fig. 1A), involves generating a large number of binder models to sample the potential interaction space. It begins with the creation of backbones for a defined topology using Monte Carlo simulations of small backbone fragments. These backbones are then docked onto a specific region of the target protein. As the binder backbones initially lack defined sequences, each docking conformation undergoes a sequence design step to obtain sequences that have a high probability of folding into the desired structure and are optimized for interaction with the target.

**Figure 1.**
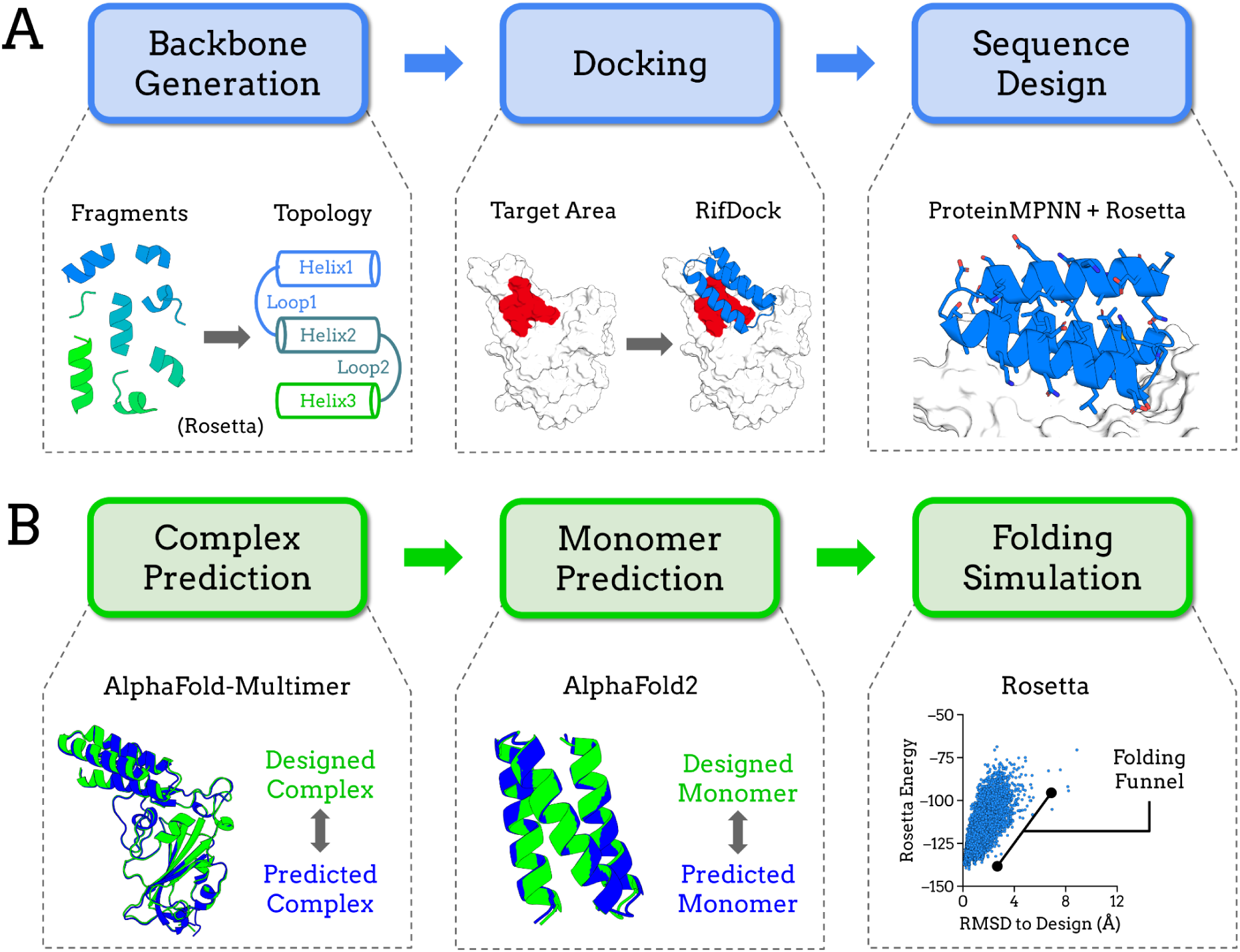
**(A)** Model Generation: the process begins with the generation of backbones with a defined topology, docked to the target region, followed by a sequence design step, where the most probable binder sequences are predicted. **(B)** Filtering: models with increased experimental viability are selected based on their structural predictions of both their complexes and monomeric forms. Folding simulations are then performed on selected binders to assess their energy landscapes.

Many models generated in this way exhibit low experimental viability. Therefore, the second phase, called *Filtering* (Fig. 1B), selects models with a higher likelihood of experimental success. This selection was based on multiple criteria, including hydrophobicity, interaction energy, and other scoring metrics, as well as structural predictions for both the complexes (target-binder) and the binder monomers. Additionally, folding simulations for selected binders were performed to analyze the energy landscape of their sequences and assess the presence of a folding funnel.

While this study focuses on CHIKV and DENV, we also applied the same protocol to the SARS-CoV-2 receptor-binding domain (RBD), a target previously used in binder design studies [35], as an additional example to demonstrate the protocol general applicability. The SARS-CoV-2 design results are in the Supplementary Material Section 10.

Each step of the protocol is described below, and additional details are available in the Supplementary Material.

### 2.1 Target Structure Preparation

X-ray crystallography-solved target structures were obtained from the Protein Data Bank (PDB). For CHIKV, the envelope protein structure (PDB ID: 3N44) [38] was used, with a resolution of 2.35 Å. For DENV, we utilized the monomer of the post-fusion trimeric envelope protein from the type 2 variant (PDB ID: 1OK8) [21], with a resolution of 2.00 Å.

As a preprocessing step, complexed proteins, solvent molecules, and linked glycans were removed from the structure. Moreover, to reduce computational cost, domains distant from the target area were removed for CHIKV and DENV. In these cases, short poly-glycine loops were created to reconnect the truncated regions, illustrated in Supplementary Material Section 1.

Following this preparation, all target protein structures were relaxed using the FastRelax [39] function from the Rosetta software (version 3.13) [40].

Additionally, pdb-tools [41] was used throughout the protocol to manipulate PDB files.

### 2.2 Backbone Generation

For CHIKV binders, we employed a three-helix bundle topology (3-helix), which was selected based on its high experimental success rate compared to other topology types [42]. In contrast, previous attempts using the 3-helix topology for DENV were ineffective, likely due to poor shape complementarity between the binder surface and the fusion loop, which has a convex, pointed geometry. We designed a custom four-helix bundle (4-helix) with tetrahedral geometry to better match the surface features of DENV and improve its interaction. This configuration allows for one or two ‘interaction pockets’ to remain accessible for interaction with the fusion loop, while preserving a stable binder structure. Representative backbones for each topology are shown in Supplementary Figure S3.

We generated backbones for each target using the Rosetta FoldArchitectMover [43], within the RosettaScripts framework [44]. Candidates were then filtered based on energetic convergence and fragment quality metrics. A total of 100 high-quality 3-helix backbones were selected for CHIKV, and 200 4-helix backbones for DENV, accounting for the increased complexity of its custom topology.

### 2.3 Docking

A higher hydrophobicity in the target area correlates with increased experimental success of designed binders [33]; therefore, a region of hydrophobic residues was defined as “target residues” for each target protein using the Spatial Aggregation Propensity (SAP) metric [45]. SAP considers the residue hydrophobicity, solvent access and other residues in proximity. The defined target areas also correspond to key sites involved in the viral entry processes: CHIKV envelope interface with MXRA8 and the DENV fusion loop, which interacts with the endosomal membrane (Fig. 2).

**Figure 2.**
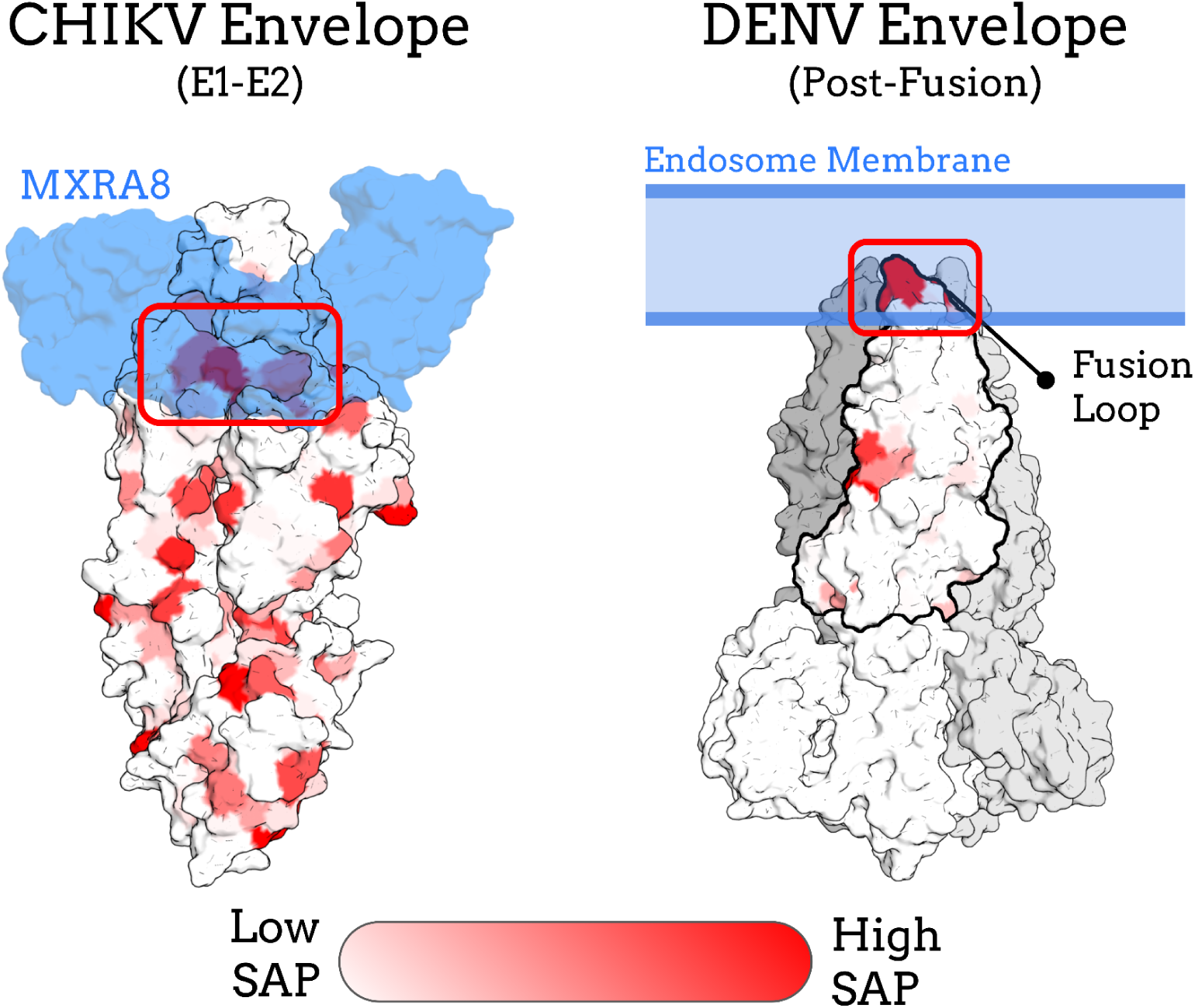
Surface hydrophobicity and target areas in the viral proteins. Target structures are colored according to their SAP score, with red indicating highly hydrophobic regions. Native binding partners are shown in blue. The target areas selected for binder design are highlighted in red boxes; they exhibit high hydrophobicity and participate in critical steps of viral entry. The DENV protein is displayed in its trimeric form, although the SAP score was calculated using the trimmed monomeric version (in bold). Structures and orientations of MXRA8 (PDB ID: 6JO8) [20] were obtained from the PDB.

The defined target residues were then passed to PatchDock [46], which generates initial conformations based on surface shape complementarity. These seeds are supplied to RifDock [47], which samples both the backbone rigid conformational space (rotation and translation) and some of the sequence space using pre-calculated interactions.

For CHIKV, the previously selected 100 three-helix backbones had 300 initial conformations generated for each in PatchDock. These were then processed with RifDock, producing 300 docking conformations per input, resulting in a total of 30,000 docking conformations per target. For DENV, considering the selected 200 four-helix backbone, the same procedure was applied and a total of 60,000 docking conformations were generated.

### 2.4 Sequence Design

To design the sequence for the binders, we used a script to cycle between sequence inference and structural optimization, employing a combination of the neural network ProteinMPNN [37] and the FastRelax [39] protocol from Rosetta, respectively, following a similar methodology described by N. R. Bennett et al. [48]. Cysteines were excluded from all designs; additionally histidines were not allowed in DENV designs to avoid significant net charge changes when transitioning from physiological pH to acidic endosomal pH. For the Rosetta structural optimization step of the DENV target, any pre-existing histidine residues were modeled in the protonated state.

### 2.5 Pre-Filtering Stage

Before applying more computationally expensive filtering methods, the binder models generated in the previous phase were initially screened using Rosetta-calculated metrics such as interaction energies, hydrophobicity, shape complementarity, net charges, and hydrogen bonds at the binding interface (detailed in the Supplementary Material, Section 5). These pre-filtering steps help eliminate low-quality candidates early on, allowing the subsequent phases to focus on more promising designs. Overall, these procedures excluded more than 90% of the initial models generated.

### 2.6 Complex Prediction

The use of deep learning-based structural prediction methods as a filtering step in binder design substantially increases experimental success rates [48]. To select models with an increased probability of experimental success, complex predictions were performed using the AlphaFold-Multimer [49] neural network, via the ColabFold [50] pipeline in version 1.3. The LocalColabFold script was used to run it locally. Multiple Sequence Alignments (MSAs) were generated using the MMseqs2 server [51] and templates were allowed. Model selection was based on the C-alpha Root Mean Square Deviation (RMSD) between the prediction and the designed structures, calculated using the TM-score software [52], as well as the AlphaFold-Multimer confidence metric (0.8 × ipTM + 0.2 × pTM).

### 2.7 Monomer Prediction

Conformational changes can occur during protein-protein interactions; consequently, the structure of a binder in its free (unbound) state, may differ from its structure in complex with the target (bound). As in small-molecule design, where minimizing entropic penalties is valuable [53], significant conformational changes or transitions from disordered to ordered states are undesirable, as they can introduce additional energetic barriers for binding. AlphaFold predictions have proven effective in identifying disordered regions [54] and context-specific conformations in protein complexes [55], [56]. Furthermore, improperly folded proteins may present challenges related to expression and susceptibility to degradation [57]. To address these concerns we evaluated the structural predictions of the binders in their monomeric states.

Monomer predictions were performed using the AlphaFold2 [36] neural network, with the ColabFold [50] pipeline in version 1.3 and executed locally with the LocalColabFold script. Although MSAs were allowed through MMseqs2, no sequences were found for any model, likely because the binders are *de novo* proteins with no relationship to natural proteins. The filtering was also based on C-alpha RMSD and confidence metrics (pLDDT).

### 2.8 Folding Simulations

As a final step, Monte Carlo fragment-based folding simulations were performed using the AbinitioRelax [58] protocol from Rosetta. Fragment libraries were generated using the automated Robetta server [59] (http://old.robetta.org/), and secondary structure predictions were obtained from the PSIPRED server [60] (http://bioinf.cs.ucl.ac.uk/psipred/). Given the relatively small size of the binders (54-60 amino acids), 10,000 decoys were generated per sequence.

Rather than simulating all designed sequences, only a subset of top candidates, ranked by interaction energy, was selected for folding simulations. For each target, simulations continued until three models exhibiting clear folding funnels [61] were identified and these were chosen for further analysis.

This final step served as a cross-validation measure, relying on a fundamentally different structure prediction methodology compared to deep learning-based approaches, such as AlphaFold. Moreover, these simulations have been shown to perform reliably for small proteins [62], [63], such as those designed in this study.

### 2.9 Molecular Dynamics Simulations

To evaluate the stability of a set of selected binder prototypes, both in complex with their targets and as isolated monomers, all-atom molecular dynamics (MD) simulations were performed using the GROMACS 2022.4 package [64]. Analyses were carried out using the GROMACS 2022.6 tools and custom Python scripts.

Simulation systems consisted of either binder–target complexes or binders free in solution, solvated using water molecules, with sufficient Na⁺ and Cl⁻ ions added to neutralize the system charge and achieve a physiological salt concentration of 0.15 mol/L. A cubic simulation box was used in all cases, with a minimum distance of 1.5 nm between any protein atom and the edge of the box.

The CHARMM36m force field [65] and the standard TIP3P water model [66] were used for all simulations. Systems were energy minimized, followed by equilibration in the canonical (NVT) ensemble for 20 ps and the isobaric-isothermal (NPT) ensemble for 100 ps, during which all heavy atoms were restrained. Production simulations were performed under NPT conditions at 300 K and 1.013 bar, using periodic boundary conditions in all directions, and a time step of 2 fs with the Leap-Frog integrator [67].

Bonds involving hydrogen atoms were constrained with the LINCS algorithm [68], and long-range electrostatic interactions were treated using the Particle-Mesh Ewald (PME) method [69]. The Verlet scheme [70], [71] was used for neighbor listing. Van der Waals and Coulomb interactions were truncated at a cut-off radius of 1.2 nm, with the *force-switch* modifier applied to van der Waals potentials starting at 1.0 nm to smoothly bring it to zero. Pressure coupling was achieved using the Parrinello-Rahman barostat [72], [73] with a coupling time constant of 10 ps and a compressibility of 4.5e-5 bar⁻¹. Temperature coupling was achieved using the V-rescale thermostat [74] with a coupling constant of 0.1 ps, treating protein molecules and the solvent (comprising water and ions) as separate groups. For each target, three binder candidates were selected (as described in the previous step) and simulated in complex with their targets and as isolated monomers, resulting in a total of 18 systems (considering the supplementary SARS-CoV-2 RBD binders). Each system was simulated for 1000 ns or 2000 ns. Additionally for DENV systems, histidines were protonated.

### 2.10 Plotting and Graphical Visualization

All plots were generated using the matplotlib package [75] in Python and all structural images were rendered in PyMOL [76].

## 3. Results and Discussion

The protocol was successfully applied to the three target proteins, and the corresponding statistics are summarized in Table 1.

**Table 1.** Number of models that passed each filtering step. Number and percentage of models progressing through each filtering step of the binder design protocol for CHIKV, and DENV targets. *Folding simulations were performed only on a subset of top candidates based on interaction energy.

| Table 1. Number of models that passed each filtering step |  |  |
| --- | --- | --- |
| Step | CHIKV | DENV |
| Model Generation | 30,000 (100%) | 60,000 (100%) |
| Pre-filtering | 1531 (5.1%) | 2037 (3.4%) |
| Complex Prediction | 100 (0.33%) | 141 (0.23%) |
| Monomer Prediction | 92 (0.31%) | 96 (0.16%) |
| Folding Simulation* | 3 out of the top 14 | 3 out of the top 25 |

More than 90% of the initial models were eliminated during the pre-filtering stage based on parameters such as interaction energy and hydrophobicity (see Section 2.5). This high elimination rate is consistent with previous studies employing classical computational design methods [9], including fragment-based backbone generation and docking. Although such extensive filtering ensures that subsequent, more computationally intensive steps are focused on the most promising candidates, it also highlights that a considerable portion of computational resources during the Model Generation phase is spent on models with low experimental viability. This is expected in classical design workflows, where the binder backbone generation occurs without explicitly incorporating the surface features of the target. In this work, efforts were made to improve this by tailoring the DENV binder backbone to accommodate the convex surface of the fusion loop; however, the surface itself was not explicitly used as an input during backbone generation.

Furthermore, the optimal binding mode or orientation for a given backbone is unknown, necessitating the generation of multiple conformations to sample the interaction landscape adequately. As a result, many of the generated models are unsuccessful; though generating a large pool increases the likelihood of finding high-quality candidates. With recent advancements of diffusion-based methods for protein backbone generation [77], some of these limitations can be addressed, as the backbone design process incorporates explicit information about the target surface and adjusts the model accordingly.

Despite these challenges, classical methods continue to deliver meaningful results for binder design, as demonstrated by the significant proportion of designs correctly predicted by AlphaFold with high confidence (Table 1).

Our findings confirmed that the concern regarding conformational differences between bound and unbound states was valid; some binders that performed well during the “Complex Prediction” did not have similar structures when predicted as monomers and were consequently filtered out. Visual inspection of some of these excluded structures (Supplementary Material, Figure S7) revealed monomeric conformations with exposed alpha-helices detached from the binder hydrophobic core, despite exhibiting high pLDDT confidence scores. Although such features may not necessarily be predictive of actual behavior in solution, they could suggest reduced experimental viability compared to correctly folded designs.

Folding simulations for CHIKV binders, which employed the 3-helix topology showed a considerable success rate, with 3 out of 14 top candidates exhibiting a folding funnel. In contrast, for the DENV binders, which were designed using a 4-helix topology, the success rate was notably lower (3 out of 25), likely due to the increased complexity of the topology.

Interestingly, a significant portion of the binders subjected to folding simulations did not display a folding funnel converging to the designed structure. Considering that these binders had previously passed two AlphaFold prediction filters with high confidence, this result may suggest that, in some cases, high-confidence AlphaFold predictions might not be enough to assert structural viability fully. This highlights the potential value of incorporating energy-based methods as complementary filters in protein design protocols.

From a practical standpoint, the application of folding simulations should be weighed against the experimental testing capacity of a given project. Since these simulations are computationally expensive and not suited to high-throughput settings, they are best suited when only a limited number of binders will be tested experimentally. Ideally, hundreds of designs are needed to reliably identify high-affinity hits [33], [48], with fewer required when using diffusion-based backbone generation methods [77]. In this work, folding simulations were used to select prototypes for subsequent evaluation using MD simulations. Alternatively, suppose high-throughput experimental methods, such as yeast display [78], are available, along with easy access to large-scale oligonucleotide acquisition. In that case, it may be preferable to skip this step and proceed directly from “Monomer Prediction” to experimental screening. As more than 99% of generated models are typically discarded in classical protein design protocols, the total number of designs produced during the initial phase should be scaled according to the target difficulty, with more challenging surfaces requiring a larger number of models generated to maximize the chances of success [33], as was done here for the DENV binders.

In the following sections, we showcase representative binders for each target, highlighting those with the most promising results. These examples represent only a small fraction of the total design pool and should therefore not be interpreted as a measure of the overall efficiency of the protocol. Instead, they serve to demonstrate the viability of the binders generated through the proposed design pipeline.

### 3.1 Complexes

The selected complexes for CHIKV (73_CHIKV), and DENV (98_DENV), along with their respective analyses, are presented below (Fig. 3).

**Figure 3.**
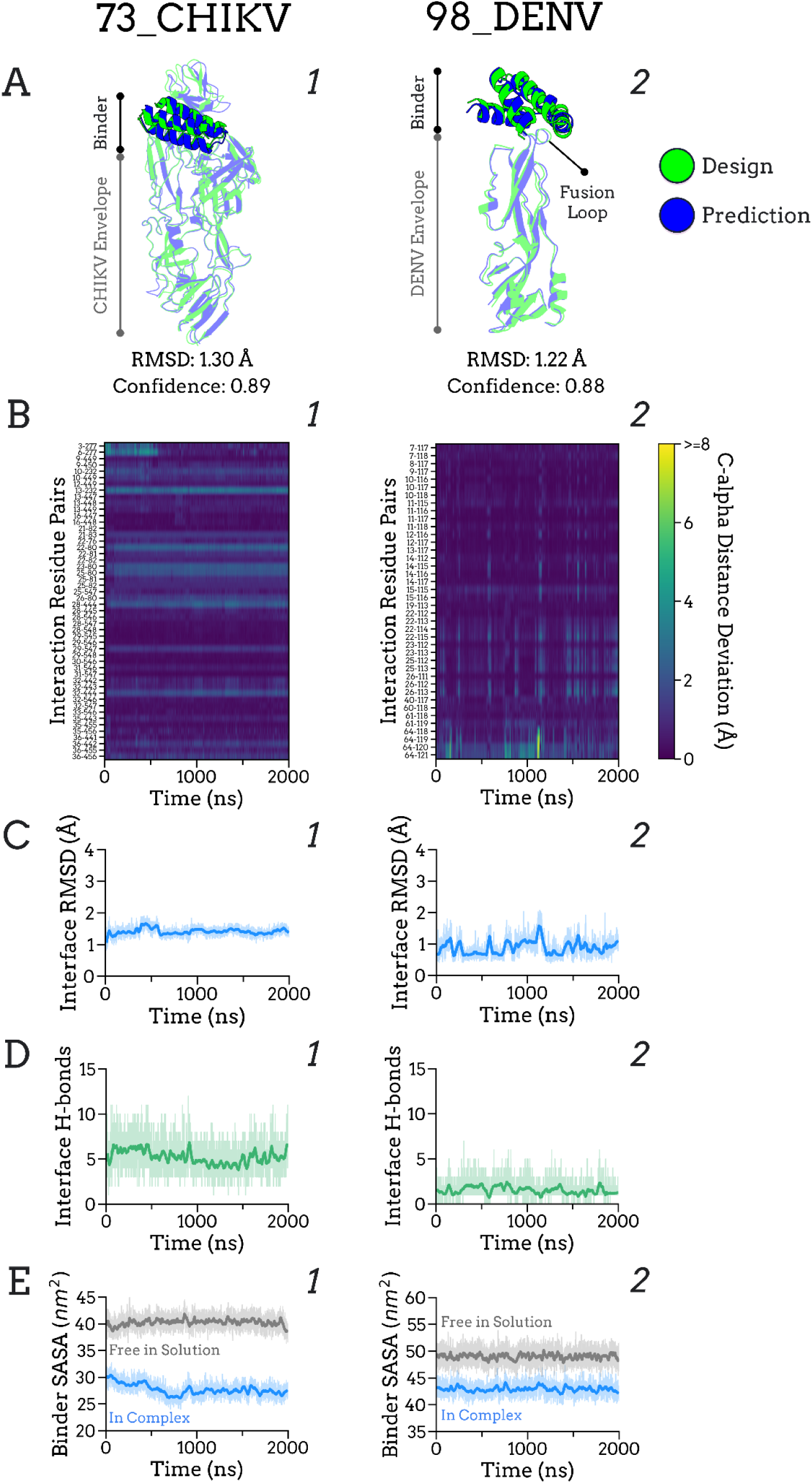
(**A**) AlphaFold-Multimer predictions of SARS-CoV-2 RBD binder complexes, with designed models (green) superimposed on predicted complexes (blue). Low RMSD values indicate high similarity between the models. (**B**) Interaction residue pair stability analysis. Each interaction pair (Cα distance < 8 Å) is plotted over time, with dark blue indicating low Cα distance deviation and yellow indicating high deviation. Y-axis labels denote residue pairs (binder residue – target residue); note that target residue numbering does not correspond to native gene numbering due to index renumbering. (**C**) Interface RMSD over time, calculated using the same interacting residues as in B. (**D**) Number of interface H-bonds over time, defined based on distance and angle criteria. (**E**) Solvent Accessible Surface Area (SASA) of the binder in complex (blue) and free in solution (grey) over time; a lower SASA in the complexed state, compared to the free state, suggests sustained interface interaction.

AlphaFold-Multimer predictions of these complexes showed high structural similarity to the original designs and present high confidence scores (Fig. 3A), indicating an increased probability of experimental success [48].

MD simulations showed stable interactions for both complexes. As illustrated by the interaction residue pair analysis (Fig. 3B), most residue pairs (Cα distance < 8 Å) showed minimal distance deviations throughout the simulations, consistent with the low and stable interface RMSD values (Fig. 3C).

The analysis of interface hydrogen bonds (H-bonds) between binder and target further supported the stability of the complexes, although with substantial fluctuations over time (Fig. 3D). A notable aspect was the low number of H-bonds in the 98_DENV complex interface, averaging approximately one H-bond throughout the simulation (Fig. 3D2). This is consistent with the small interface of the DENV fusion loop, primarily composed of three hydrophobic residues: Trp112, Leu118, and Phe119 (native numbering: Trp101, Leu107, Phe108), and a limited region of exposed backbone atoms.

A complementary analysis involved calculating the Solvent Accessible Surface Area (SASA) of the binders in both complexed and free states (based on monomer simulation data). This is based on the rationale that, in stable protein–protein interactions, the binding surface of the binder remains buried and shielded from solvent exposure, resulting in a lower SASA when in complex. If a dissociation event were to occur, the binder’s SASA would increase to values similar to the free (unbound) state. For both binders, the complexed SASA values remained consistently lower than those in the free state (Fig. 3E), indicating a consistent interaction throughout the simulations. In the case of 98_DENV, the SASA difference between bound and unbound was modest (Fig. 3E2), reflecting its relatively small interaction interface.

Overall, these results indicate that the proposed protocol successfully generated binder models with viable interaction interfaces with their respective targets.

### 3.2 Monomers

In addition to the complex analyses, the selected binders were also evaluated as monomers, free in solution, to assess their structural stability in the absence of the target (Fig. 4).

**Figure 4.**
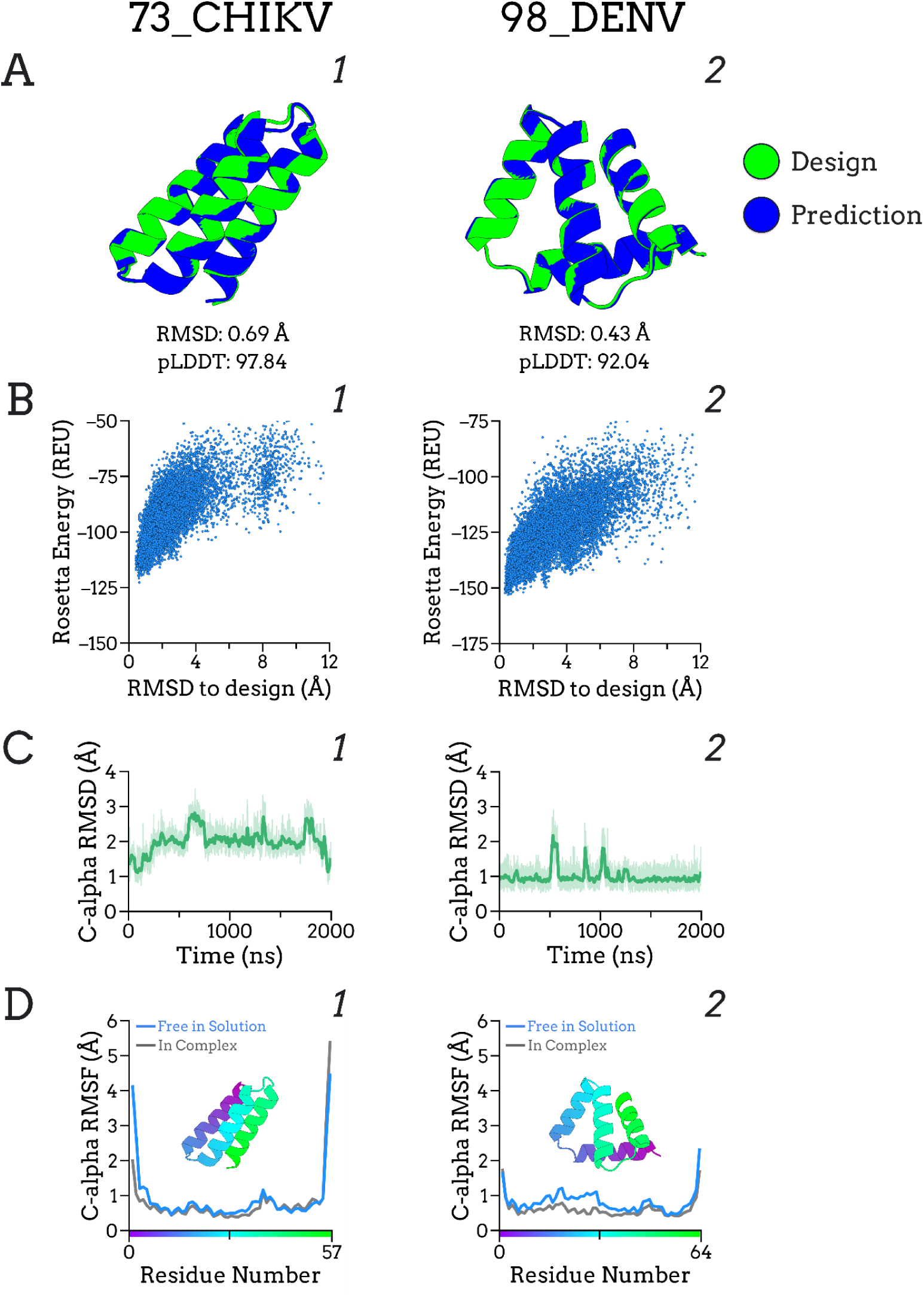
Monomer predictions, folding simulations and MD simulation analysis. (**A**) AlphaFold2 predictions for the three selected binders (73_CHIKV and 98_DENV) as monomers. Designed models (green) are superimposed onto the predicted structures (blue). The low RMSD indicates high similarity between the models. (**B**) Folding Simulations. Energy versus RMSD plots show a folding funnel, with decoys converging to the reference structure (the designed model). (**C**) Binder Cα RMSD over time during MD simulations. (**D**) Binder Cα RMSF in MD simulations, both free in solution (blue) and in complex (grey). The X-axis corresponds to the residue numbering, which is also reflected in the structural coloring.

AlphaFold2 predictions showed high structural similarity to the designed models and present high pLDDT (Fig. 4A). At the same time, Rosetta folding simulations revealed folding funnels with models converging toward the reference structure (Fig. 4B), suggesting a favorable propensity for the binder to adopt the intended fold.

MD simulations further supported the overall stability of the binder structures, indicated by the low RMSD overtime (Fig. 4C). Minor increases and spikes in RMSD were attributed to flexibility at the C and N-terminals, as observed in the RMSF plots (Fig. 4D). Importantly, the flexibility of the binders did not differ substantially between the complexed and free states, suggesting that the binding event may not require extensive transitions from disordered to ordered conformations, which could have a positive effect in the interaction (see Section 2.7).

The binder 73_CHIKV exhibited a noticeable decrease in RMSF for its N-terminal residues when in complex (Fig. 4D1), likely due to their proximity to the interaction interface. Visual inspection revealed that these residues are positioned at the periphery of the interface, rather than being deeply buried, where they likely form interactions with the CHIKV envelope (Supplementary Material, Figure S8). This observation suggests that these residues are highly flexible when the binder is free in solution, and that interaction with the target promotes local stabilization.

Collectively, these results suggest that the protocol successfully balances the two key aspects of binder design, interface optimization and monomer foldability, producing binders with viable interaction interfaces while maintaining structural stability in the bound and unbound states

It is important to note that the binders showcased here represent the most promising cases. Some additional systems (see Supplementary Material, Section 11), did not exhibit similar structural or interaction stability during MD simulations. This is expected, given that *de novo* protein design protocols often have relatively low experimental success rates. Accordingly, the presence of unstable binders should not be viewed as a limitation of the protocol itself, but rather as an inherent challenge of the task and an ongoing need to improve design, filtering, and selection strategies.

## 4. Conclusion

To our knowledge, this work represents the first systematic generation and evaluation of de novo designed miniprotein binders specifically targeting critical entry proteins of Chikungunya and Dengue viruses, two neglected tropical diseases. In addition, we present a generalizable protein design protocol with detailed rationale for each step, to aid the community in addressing current and future challenges of similar nature. The challenge of poor shape complementarity faced in designing DENV binders, and how it was overcome, offers a conceptual framework for addressing similar geometric constraints in future protein design efforts.

While the protocol relies on classical backbone design methods that require extensive sampling and the generation of many models, most of which are discarded, it nonetheless succeeded in producing binders with viable interaction interfaces and stable monomeric structures. The emergence of diffusion-based backbone design offers a promising approach to improve efficiency by explicitly incorporating target surface information [77]. As the field of protein design evolves rapidly, the ongoing update and integration of new methodologies will be crucial for maintaining and enhancing the effectiveness of future design pipelines.

The binder models generated in this work represent potential biopharmaceuticals for the treatment of CHIKV, and DENV infections, serving as a foundation for further optimization. However, experimental validation remains essential to confirm biological activity and to refine the design protocol with empirical data.

## 5. Data Availability

The initial backbones, binders analyzed, and scripts used in this study are available at the GitHub repository: https://github.com/danilokiyoshi/SM_Binder_Design_CHIKV_DENV.

## 6. Acknowledgments

This work was supported by the National Council for Scientific and Technological Development (CNPq - 465259/2014-6; CNPq - 304155/2021-7), the Coordination for the Improvement of Higher Education Personnel (CAPES), the National Institute of Science and Technology Complex Fluids (INCT-FCx) and the São Paulo Research Foundation (FAPESP - 2014/50983-3; FAPESP - 2023/09291-0; FAPESP - 2024/14345-4). Computational resources were provided by the National Laboratory of Scientific Computation (LNCC - 224709).

