## Supplementary Material for "Computational Design of Miniprotein Binders Targeting the Viral Entry in Chikungunya and Dengue, Neglected Tropical Diseases"

Danilo Kiyoshi Matsubara[1], Peter Park[2,3], Iolanda Midea Cuccovia[1], Hernan Chaimovich[1]

[1] Department of Biochemistry, Institute of Chemistry, University of São Paulo, Brazil.

[2] Department of Clinical and Toxicological Analyses, Faculty of Pharmaceutical Sciences, University of São Paulo, Brazil.

[3] Present address: Department of Biological Sciences, College of Arts and Sciences, Lehigh University, USA.

Danilo Kiyoshi Matsubara,

Peter Park,

Iolanda Midea Cuccovia,

Hernan Chaimovich,

Keywords: Computational protein design, Antiviral binder design, Chikungunya virus (CHIKV), Dengue virus (DENV), Molecular dynamics (MD) simulations, Neglected tropical diseases, Viral entry inhibition.

### METHODS

Complementary information about the methods is displayed here. It’s intended to be read together with the main text, as key informations are not repeated. The SARS-CoV-2 RBD binder methods are also included.

#### Section 1: Target Structure Preparation

##### CHIKV

The CHIKV envelope structure used in this work was composed of the three envelope proteins E1, E2 and E3. Since our target surface is located on the interface of E1 and E2, the E3 chain was removed.

To reduce computational costs, protein domains of E1 distant from the target region were removed, and the resulting construct was connected using a short glycine loop to preserve structural integrity (Fig. S1A). AlphaFold-Multimer [1] predictions of this partial E1-E2 complex showed concordance to the intended structure, suggesting the trimming should not affect the target area, at least in the “Complex Prediction" step of the protocol. As such, this partial E1-E2 was used as target protein in the protocol.

Using the previously generated MD simulations of the 73_CHIKV complex, we examined whether the target (partial envelope) showed any signs of unfolding. No significant unfolding was observed (Fig. S1B). As expected, RMSF values were elevated for the glycine loop, but were comparable to those of other flexible loops within the structure. It’s important to note that the data might be influenced by the binder which is complexed to the target.

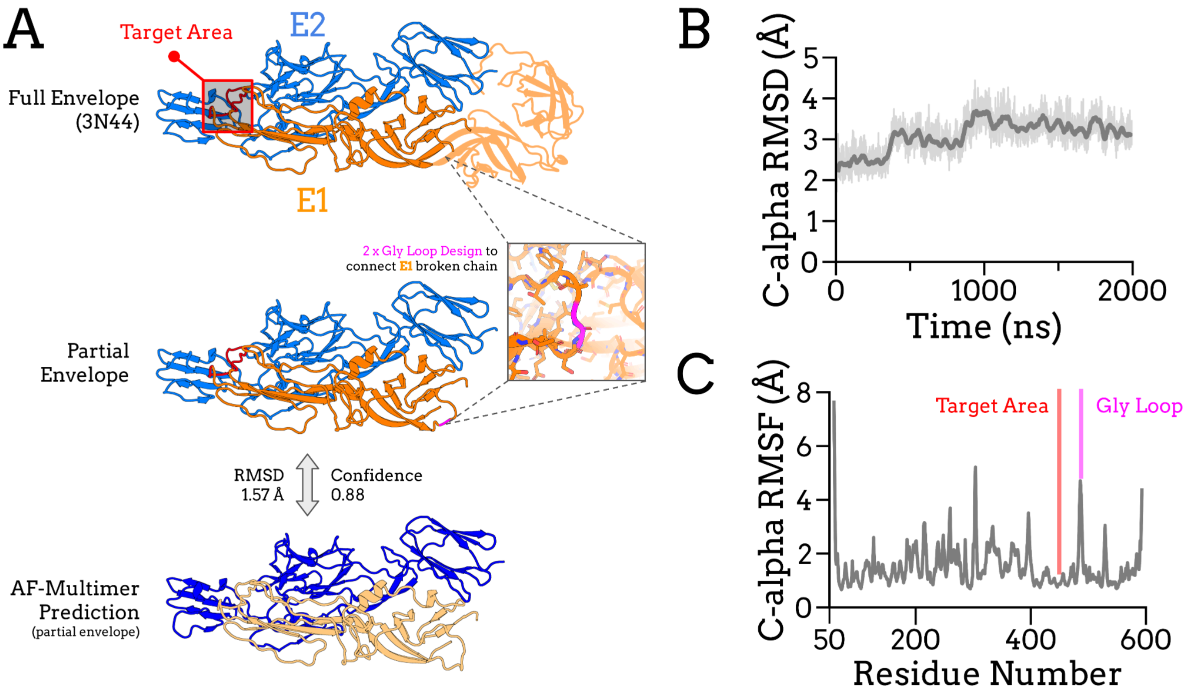


**Figure S1.** **(A)** CHIKV envelope trimming process. The full envelope consists of the proteins E1 (orange) and E2 (light blue). Domains removed from the construct are depicted with transparency. The designed glycine loop reconnecting the remaining E1 protein is shown in magenta. The target area is colored in red The final construct used in this study is referred to as the 'Partial Envelope.' AlphaFold-Multimer predictions were employed to qualitatively assess the structural effects of domain removal. Confidence was calculated as 0.8 × ipTM + 0.2 × pTM. **(B)** C-alpha RMSD of the CHIKV_73 target (partial envelope) shows stable values, indicating overall structural stability. **(C)** C-alpha RMSF of the target (partial envelope), with the glycine loop highlighted in pink and the target region in red. *MD simulations were performed with the target complexed to the binder, so potential effects of the binder on the target cannot be excluded.

The postfusion envelope protein of DENV was utilized in this study. Given its trimeric nature, only one monomer was used in the protocol. Similarly to the CHIKV envelope, we trimmed off domains distant from the fusion loop (target area) to decrease the computational cost. AlphaFold2 [2] predictions were conducted on the partial envelope to assess its alignment with the intended construct. In MD simulations of the complex 98_DENV, unfolding of the target has not been found (Fig. S2). As with the previous target analysis, the data for this case were obtained in complex with the DENV binder, which may influence the results.

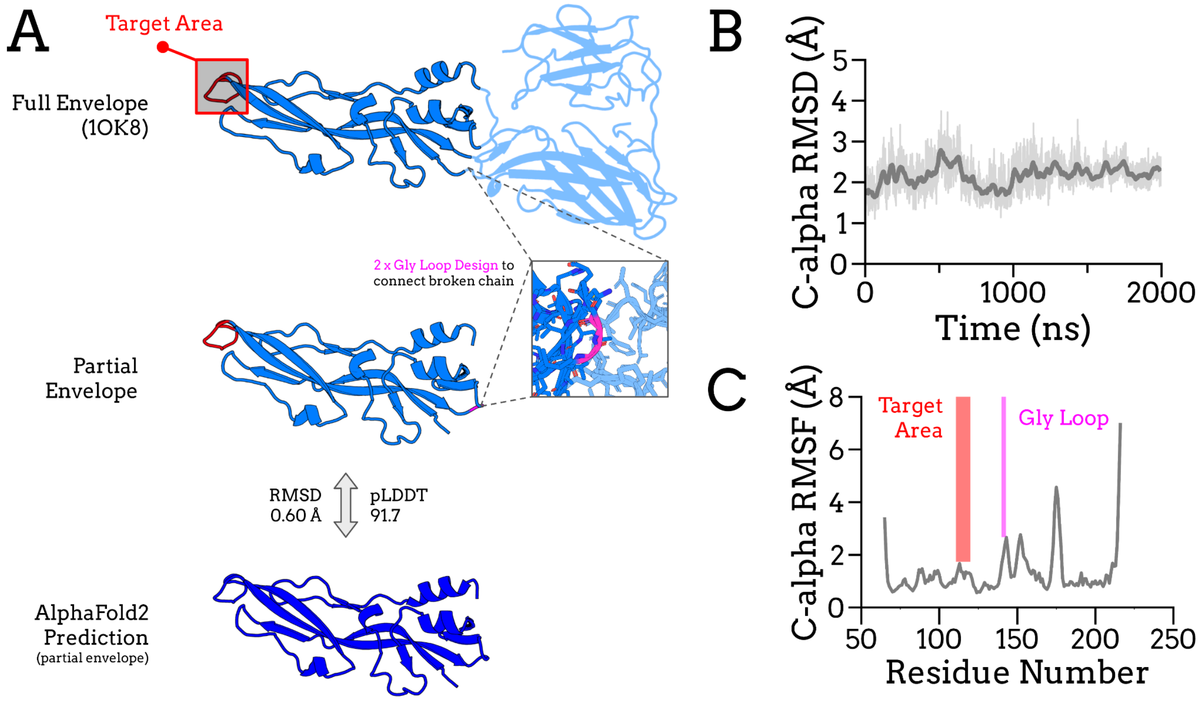


**Figure S2. (A)** DENV envelope trimming process. One monomer is presented in light blue. Domains removed from the construct are depicted with transparency. The designed glycine loop reconnecting the remaining domain is shown in magenta. The target area is colored in red. The final construct is referred to as the 'Partial Envelope.' AlphaFold2 predictions were employed to qualitatively assess the structural effects of domain removal. **(B)** C-alpha RMSD of the 98_DENV target (partial envelope). **(C)** C-alpha RMSF of the target (partial envelope), with the glycine loop highlighted in pink and the target region in red. *MD simulations were performed with the target complexed to the binder, so potential effects of the binder on the target cannot be excluded.

##### RBD

The RBD of SARS-CoV-2 (PDB ID: 7ZXU) [3] has a resolution of 1.89 Å and belongs to the BA.4/BA.5 Omicron variant.

As a cleaning step, neutralizing proteins, solvent molecules and the linked glycan group (Asn343) were removed from the structure. The glycan group was excluded because it cannot be modeled by some tools within the protocol, and it is unlikely to affect the design process due to its distance from the target area.

**Section 2: Backbone Generation**

For SARS-CoV-2 RBD and CHIKV, the chosen topology for the binder was a three helical bundle (3-helix), selected due to its high experimental success rate compared to other topologies [4].

This topology was then defined using the FoldArchitectMover [5] function from Rosetta [6], within the RosettaScripts [7] framework, as follows:

- Three α-helices (H1, H2, and H3), each comprising 15–17 residues.
- Two loops (L1 and L2), each with 3–4 residues, connecting H1 to H2 and H2 to H3.
- Helix pairing was defined as: H1–H2 antiparallel, H1–H3 parallel, and H2–H3 antiparallel.

The pairing criteria for helices was defined as distance between helix centers of mass < 12.5 Å and inter-helix angle < 20°. During the project execution it was noticed that using the energy function default values resulted in the generating of poorly compact backbones, which could negatively affect the next steps of the protocol. To address this, the contribution of the radius of gyration (rg) term within the energy function was multiplied by 2 to promote more compact conformations.

A total of 2,000 backbones were generated using the parameters described above. Of these, 744 passed the internal energetic convergence and internal filtering of FoldArchitectMover. Further selection was performed using fragment quality metrics, specifically worst9mer (< 0.4 Å) and worst9mer_helix (< 0.15 Å). These metrics assess structural similarity to natural fragments based on RMSD: worst9mer refers to the single 9-residue backbone fragment of the structure with the greatest deviation from the general fragment library, while worst9mer_helix considers only helical regions [4]. The lower the value for these metrics, the more “natural” the backbone is, meaning it's closer to the natural fragment database. Based on these criteria, 369 high-quality backbones were retained, and the top 100 were selected, ranked by worst9mer (the lower the better), for the next stage of the protocol.

For DENV, previous design attempts using the 3-helix topology were proven to be ineffective (no successes in AlphaFold predictions), likely due to poor shape complementarity between the binder surface and the fusion loop, which presents a convex, pointed geometry. To target the DENV fusion loop, we designed a customized topology with improved shape complementarity to its pointed surface. Specifically, a four helical bundle (4-helix) with a tetrahedral arrangement was idealized.

Also using FoldArchitectMover, this topology was defined as:

- Four helices (H1, H2, H3 and H4), each of which can have 13 or 14 residues.
- Three loops (L1, L2 and L3), L1 and L3 can have 3 or 4 residues, L2 only 1 residue. Each one connects the previously mentioned helices.
- H3 and H4 are paired in an parallel orientation

Helix pairing was defined by a center-of-mass distance < 13 Å and an inter-helix angle < 30°. Distance constraints are also added to guide the fragment simulation. The contribution of the radius of gyration (rg) term within the energy function was multiplied by 2.5.

A total of 2,000 backbones were again generated. Following convergence and internal filters, 1,886 models were retained. After applying fragment quality filters, worst9mer (< 0.4 Å) and worst9mer_helix (< 0.15 Å), 521 high-quality candidates remained, and the top 200 were selected, ranked by worst9mer, for use in the protocol for DENV.

Backbones generated here are only composed of valine residues.

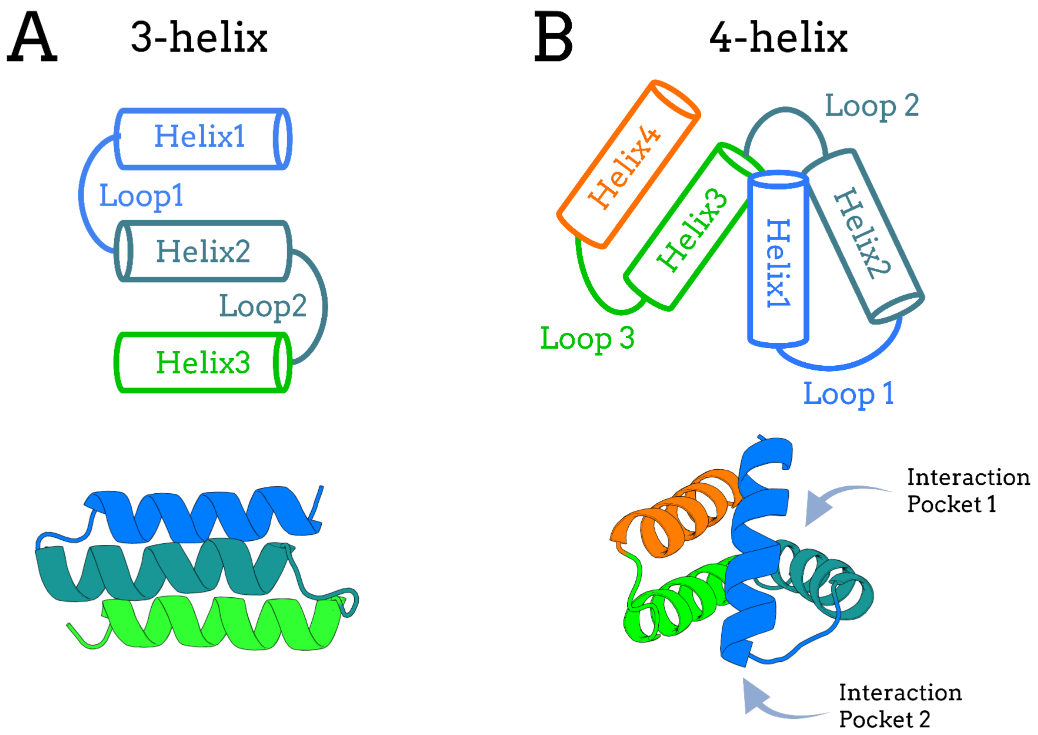


**Figure S3.** Representative topologies and structures of the (**A**) 3-helix and (**B**) 4-helix backbones. Each backbone segment is colored according to the topology shown above. Potential interaction pockets are indicated in the 4-helix backbone. Since a wide variety of conformations are sampled during backbone generation, the final models may differ from the representative examples displayed here.

#### Section 3: Target Areas and Docking

At this step, the binder backbones are composed of valines and must be docked to the target interface, after which the sequence is subsequently optimized for target interaction. To define the target area, the Spatial Aggregation Propensity (SAP) [8], was calculated with the Rosetta software, using the PerResidueSapScoreMetric function in RosettaScripts. A patch of 3-5 residues with high SAP are then chosen as target area and passed to PatchDock [9]. Protein structures colored by SAP score were made using a script available at the Supplementary Material of Cao et al [10] together with PyMol.

As for the second docking software, RifDock [11], it's actually composed of two phases: the first one, RifGen, samples millions of disembodied side chain conformations in the target region, considering their atomic interactions with the target residues, creating an "interaction field" of residues. Subsequently, in the second phase, RifDock, the backbones are docked to these precomputed side chains (Fig. S4). A practical guide to its use is available at the Supplementary Material of Cao et al [10].

To use RifDock a Rosetta 3.9 version was needed.

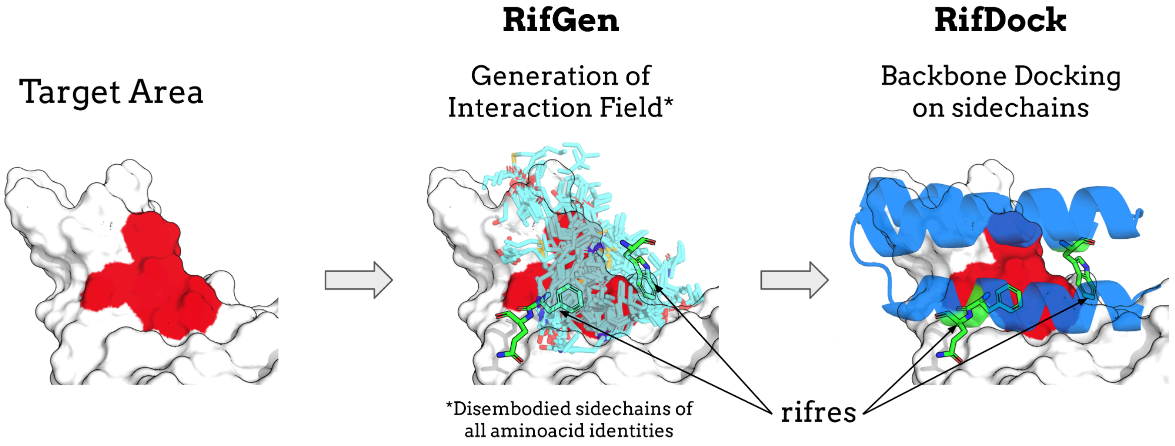


**Figure S4.** RifDock and RifGen. Given a target area, billions of possible interactions between side chains and target residues are generated (RifGen), backbones are then docked to these side chains, considering favorable geometries and conformations (RifDock). The side chains selected to be part of the docked backbone are called rifres.

The use of this type of docking methodology is necessary considering the volume at the interaction interface after docking the backbones, consisting only of valines. Conventional docking methodologies would use these valines as interaction points directly. This could result in a lack of adequate spacing in certain areas of the interface to accommodate possible amino acids with a larger volume than valine, such as tryptophan or arginine, during the sequence design process. Favorable interactions could be missed due to a lack of sufficient volume for possible substitutions. When performing docking by RifDock, the C-alphas of the backbones are used as interaction points with pre-calculated side chains. The obtained interactions (rifres) may not be ideal, but they are sufficient to obtain a more adequate available volume at the interface for the subsequent sequence design process. The adoption of flexible backbones during sequence design (adopted in the next step) can also help alleviate the issue of accommodating optimized residues at the interface.

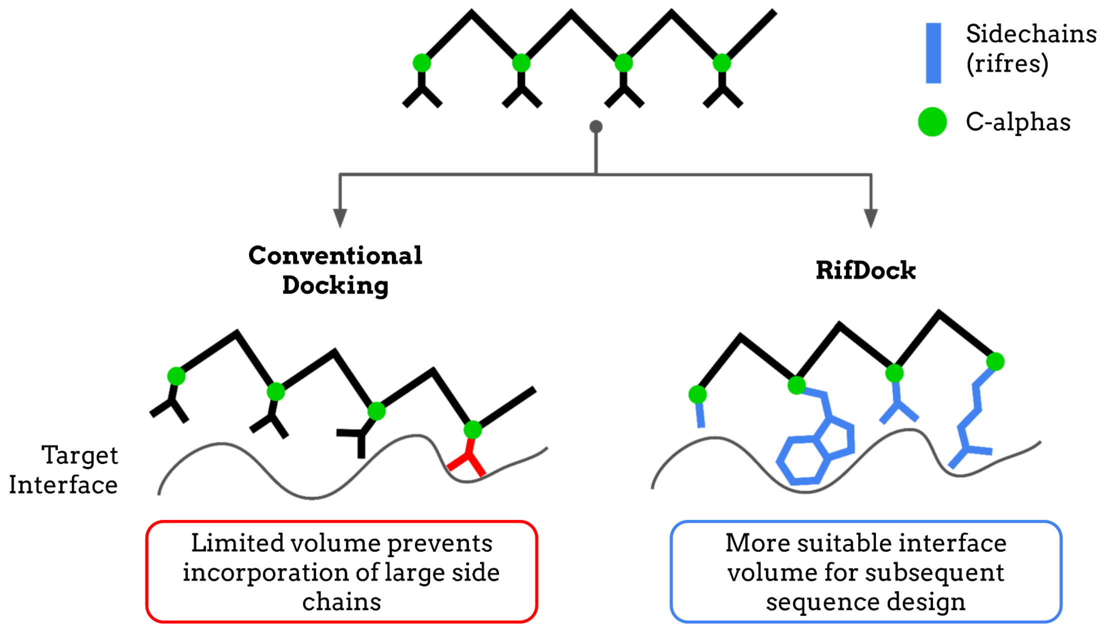


**Figure S5.** Comparison between Conventional Docking and RifDock. In conventional docking (left), valine backbone residues serve as direct interaction points, which can lead to conformations with insufficient volume (red) to accommodate side chains larger than valine. In RifDock (right), Cα atoms (green) are used as interaction points with precomputed side-chain conformations at the target interface (blue, rifres). This approach preserves a more suitable volume at the interface for subsequent sequence design.

**Section 4: Sequence Design**

A negative bias of -0.5 for lysine was applied in the ProteinMPNN [12] neural network to favor negatively charged miniproteins as it was experimentally seen that a negative net charge increased the likelihood of experimental success in binder design [4]. The bias value may need to be adjusted according to the target topology and binder size. A negative bias was applied to lysine rather than arginine, as the ProteinMPNN model shows a tendency to introduce more lysines than arginines [12].

The ProteinMPNN model used was "*v_48_030*", trained with a noise level of 0.3 Å. This choice was based on the ProteinMPNN publication [12], which reported that introducing this level of noise increases the likelihood of generating sequences correctly predicted by AlphaFold. The sampling temperature (*sampling_temp*) was set to 0.1.

For conformational optimization with the *FastRelax* protocol, the backbones of both target and binder were treated as flexible. Specific weights (*relaxscript*) for protein-protein interfaces called "*InterfaceRelax2019*" were used; it controls how the Lennard-Jones potential is varied throughout the optimization. The energy function applied was *beta_nov16*.

Five cycles of ProteinMPNN and Rosetta FastRelax [13] were applied to each docking conformation.

**Section 5: Pre-filtering**

Many models generated in the previous stage exhibit features that reduce their likelihood of experimental success, such as low target contact, high hydrophobicity, extremely positive or negative net charge, or near-neutral charge. To eliminate such candidates, a filtering step was applied to all binders, using Rosetta, based on the following metrics:

- Interaction Energy (dG_separated):
  This metric reflects the difference in energy (in Rosetta Energy Units, REU) between the bound and unbound states of the complex. Models with interaction energies higher than -30 REU were discarded, following the threshold used by Cao et al. [10].

- Binder Hydrophobicity (sap_score):
  Highly hydrophobic binders are prone to aggregation. The SAP score quantifies surface hydrophobicity and is calculated for the binder free in solution. Binders with SAP scores above 35 were removed, following the threshold used by Cao et al. [10].

- Hydrophobic Contribution at the Interface (binder_delta_sap):
  This metric quantifies the hydrophobic contribution to binding, calculated as the difference in SAP score between the unbound and bound states of the binder. The SAP depends on the solvent-accessible surface area, which is reduced upon interaction at the binding interface. A greater reduction in SAP reflects a stronger hydrophobic component of the binder's interaction interface. Since the hydrophobic effect is fundamental for binding [10] this metric allows for the elimination of models with highly polar interaction

interfaces (with low hydrophobicity). Binders with deltaSAP less than 12 were excluded, following the threshold used by Cao et al. [10].

- Interface Packing (contact_molecular_surface):

  This metric measures shape complementarity at the binder–target interface. Voids and unfilled regions decrease this value. It is proportional to the interaction surface area. Given that the 3-helix backbones used in this work for CHIKV and SARS-CoV-2 binders are similar to those in Cao et al. [10], the same cutoff of 450 was applied. For DENV binders, which have a smaller surface area, a cutoff of 320 was used. Models scoring below these thresholds were discarded. For backbones with substantially different topologies, the threshold may need to be adjusted.

- Net Charge (net_charge):

  Experimental success rates tend to be higher for negatively charged binders [4]. Since ProteinMPNN does not allow explicit control over net charge during sequence generation, a negative bias against lysine residues was introduced (see Supplementary Material, Section 4) to favor negatively charged designs. A range of allowed net charges was established; binders with net charges above -5 or below -15 were discarded, based on empirical success probabilities [4].

- Unsatisfied Hydrogen Bonds at the Interface (delta_unsatHbonds):

This metric quantifies the number of unsatisfied hydrogen bonds at the interaction interface. An unsatisfied hydrogen bond is defined as an atom capable of forming a hydrogen bond but not engaged in one [14]. The presence of unpaired hydrophilic residues at the interface is one example that increases this metric. In this work, a relatively high number of unsatisfied interface hydrogen bonds was observed; therefore, as a precaution, a maximum threshold of 9 unsatisfied hydrogen bonds at the interface was applied.

**Section 6: Complex Prediction**

The number of recycles was kept at the default value of three, and the neural network model employed was "*alphafold2-multimer-v2"*. Multiple sequence alignments (MSAs) were generated using the "UniRef+Environmental" database with the "unpaired+paired" pairing mode. The predicted models were further refined in Rosetta using the *FastRelax* protocol [13] with Cα restraints.

The confidence metric was calculated as described in the AlphaFold-Multimer methodology [1]: 0.8 × ipTM + 0.2 × pTM.

For DENV and SARS-CoV-2, predicted models with a confidence value below 0.8 and a Cα RMSD to the designed model greater than 1.5 Å were discarded. During the project, we observed that some CHIKV binder models passed the 0.8 confidence cutoff, yet the predicted structures did not exhibit tight binder–target interactions visually. We hypothesized that the elevated confidence scores were inflated by interactions between E1 and E2. To address this, we applied a stricter cutoff of 0.86 for CHIKV, which resulted in better filtered models.

**Section 7: Monomer Prediction**

The number of recycles was kept at the default value of three, and the neural network model employed was “*alphafold2_ptm*”. Multiple sequence alignments (MSAs) were generated using the “UniRef+Environmental” database with the “unpaired+paired” pairing mode. However, as mentioned in the main text, no MSAs could be successfully constructed. The predicted models were further refined in Rosetta using the *FastRelax* protocol with Cα restraints.

The reference structure of the binder, used for comparison with the predicted models, was taken from the filtered complexes of the previous step. The reference monomers were structurally optimized in Rosetta using the unconstrained *FastRelax* protocol prior to comparison, to relax minor conformational changes induced by interaction with the target.

In this step, the confidence metric used was the average pLDDT of the structure. Predicted models with a pLDDT below 90 or a C-alpha RMSD to the designed model greater than 1.2 Å were discarded.

**Section 8: Folding Simulations**

The Rosetta AbinitionRelax protocol [15] utilizes Monte Carlo simulations to explore the protein’s conformational space. To reduce the degrees of freedom, fragments of 3 and 9 residues, extracted from experimentally solved protein structures, are used to represent local chain conformations. The simulation can be understood as an exploration of the sequence’s energy landscape, assuming that the reference structure corresponds to the global minimum.

Each model generated in the simulation has an associated energy (Rosetta Energy Unit) and an RMSD relative to the reference structure. An energy versus RMSD plot is then generated, allowing the assessment of whether a folding funnel [16] was obtained, with models converging toward the reference structure (Fig. S6), this would support the idea that the sequence could adopt the designed/ideal reference structure.

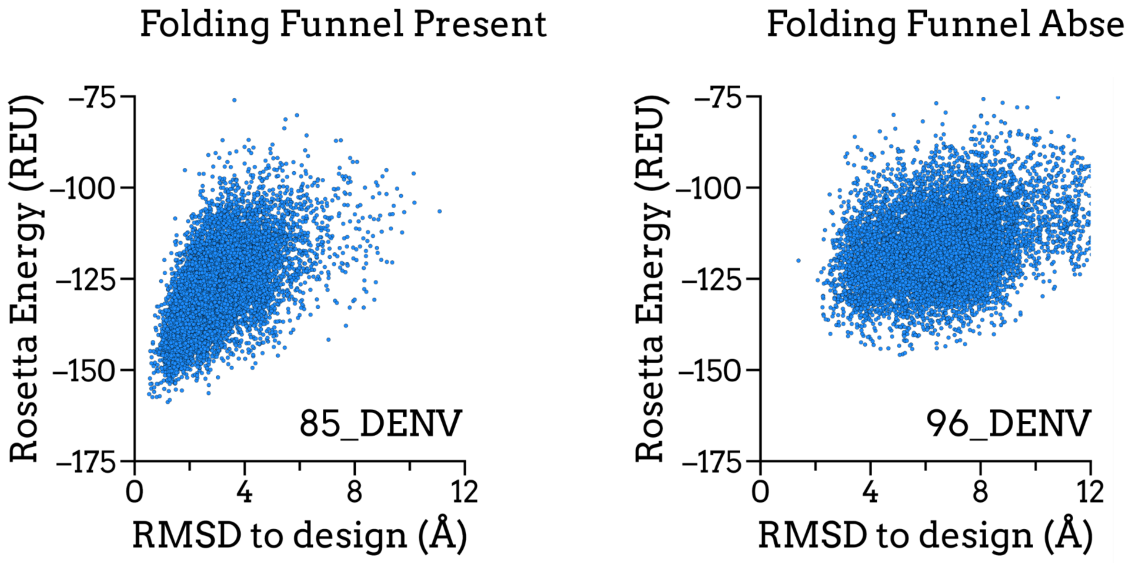


**Figure S6.** Examples of Rosetta Folding Simulations. Energy versus RMSD plots are shown, with each point representing a model (structure) obtained from an independent Monte Carlo simulation. On the left, a folding funnel can be observed, with models converging toward zero RMSD (reference structure), supporting folding to the target structure. On the right, no clear folding funnel is observed, indicating non-ideal folding relative to the reference structure and suggesting that the reference structure may not represent the global energy minimum.

#### Section 9: Complementary Figures to Main Text

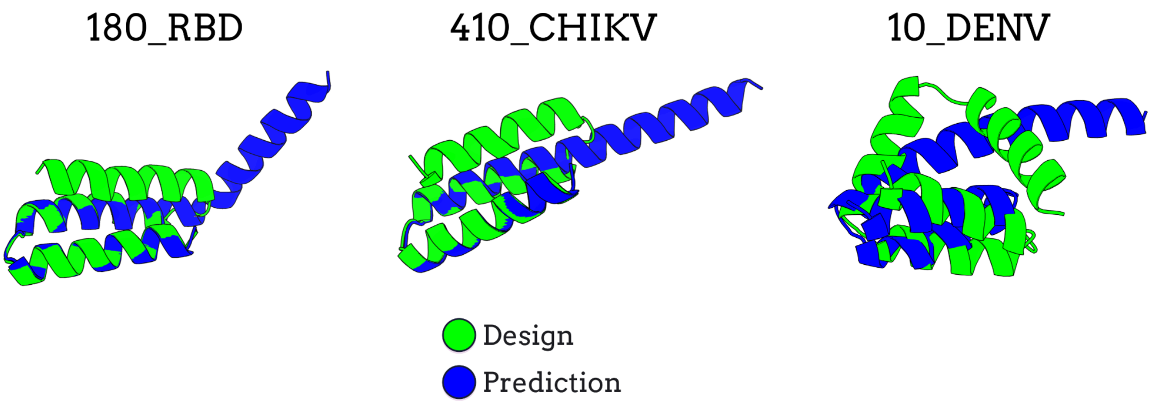


**Figure S7.** Example of Failed designs in Monomer Prediction step of the protocol. Structural alignments between designed models (green) and predicted models (blue) are displayed.

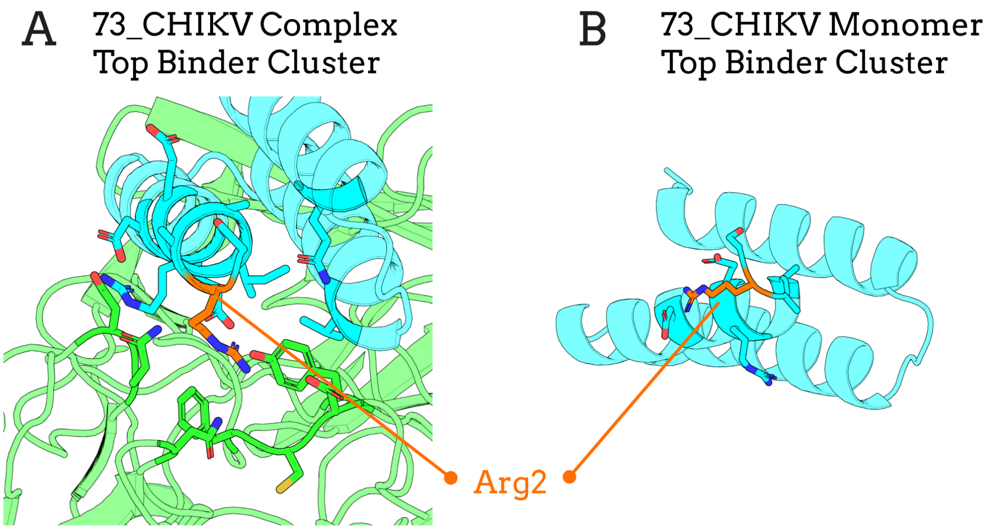


**Figure S8.** Top cluster conformation of the 73_CHIKV binder in complex (A) and free in solution (B). In the complex, Arg2 primarily interacts with the CHIKV envelope. When free in solution, Arg2 appears flexible and forms transient interactions with nearby negatively charged residues.

#### Section 10: Supplementary SARS-CoV-2 RBD Binders

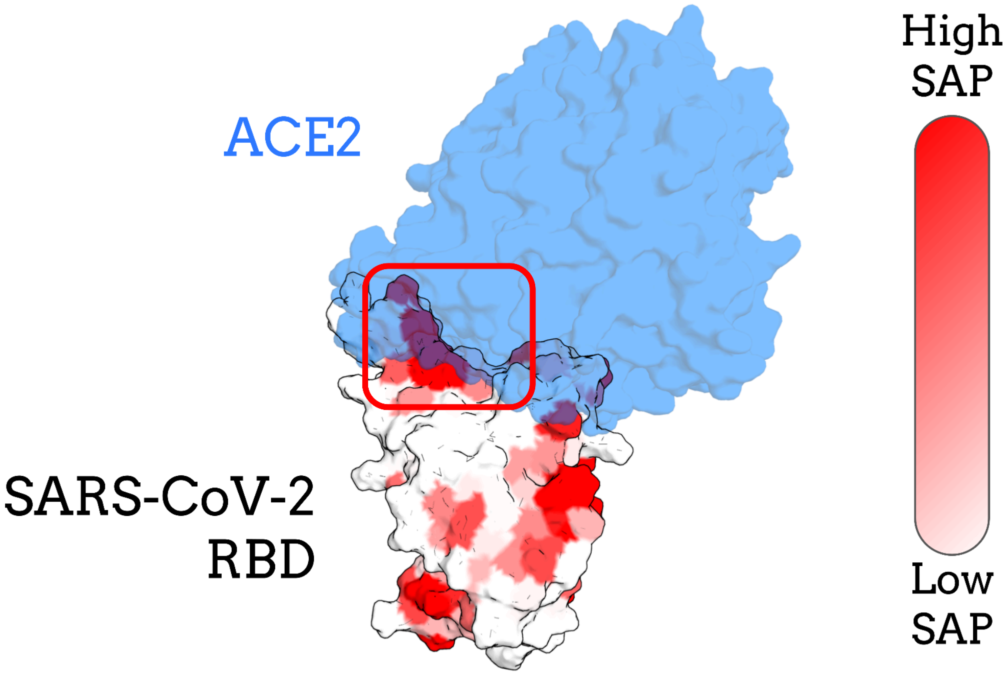


**Figure S9.** Surface hydrophobicity and target areas in the SARS-CoV RBD. The target structure is colored by SAP score, with red indicating highly hydrophobic regions. The native binding partner, ACE2, is shown in transparent blue. The target area selected for binder design is highlighted with a red box; this region participates in key interactions with ACE2 during SARS-CoV-2 viral entry. The ACE2 structure and orientation were taken from the PDB (ID: 6VW1) [17].

| **Supplementary Table 1. Number of models that passed each filtering step** | |
|---|---|
| Step | SARS-CoV-2 RBD |
| Model Generation | 30,000 (100%) |
| Pre-filtering | 1345 (4.5%) |
| Complex Prediction | 137 (0.46%) |
| Monomer Prediction | 120 (0.40%) |
| Folding Simulation* | 3 out of the top 9 |

**Table S1.** Number and percentage of models progressing through each filtering step of the binder design protocol for the SARS-CoV-2 target. *Folding simulations were performed only on a subset of top candidates based on interaction energy.

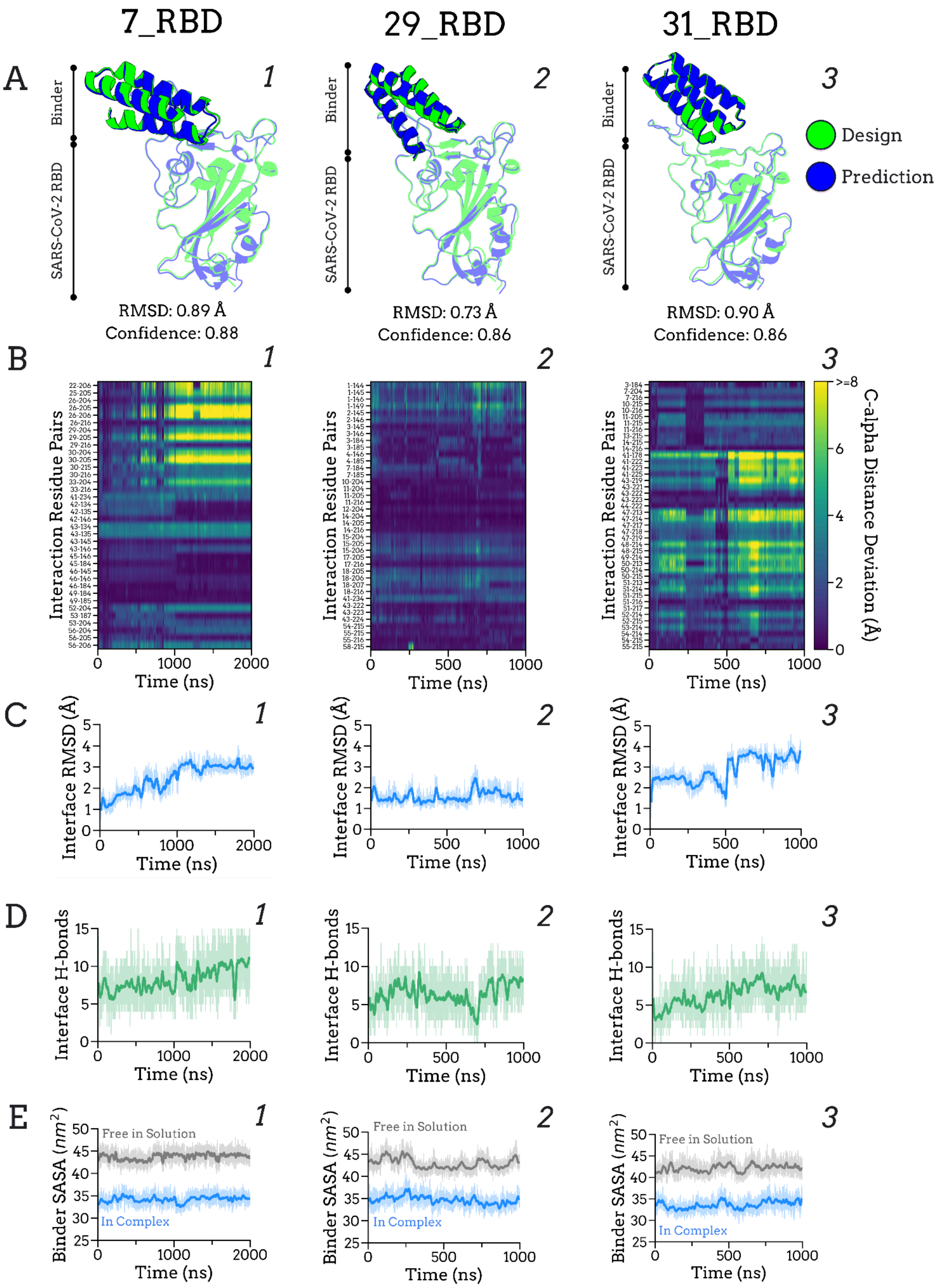


**Figure S10.** Analysis of supplementary SARS-CoV-2 RBD binder complexes. (**A**) AlphaFold-Multimer predictions with designed models (green) superimposed onto predicted complexes (blue). (**B**) Interaction residue pair stability analysis. (**C**) Interface RMSD over time. (**D**) Number of interface H-bonds over time. (**E**) Solvent Accessible Surface Area (SASA) of the binder in complex (blue) and free in solution (grey) over time. The 7_RBD system was simulated for 2000 ns, whereas all other systems were simulated for 1000 ns.

Taking 7_RBD as a representative binder for SARS-CoV-2, it exhibited a gradual increase in interface RMSD during the first 1000 ns before reaching stability (Fig. S10, C1). This behavior appears to involve residues 204–206 (native numbering: 475–477) of the RBD. Visual inspection (Fig. S12) confirmed that these residues belong to a flexible loop, which temporarily dissociates from the interface. Despite this local flexibility, 7_RBD maintained overall interaction stability, likely supported by other residue pairs (Fig. S10, B1). Additionally, 7_RBD displayed a gradual increase in the number of interface H-bonds over time (Fig. S10, D1), suggesting a possible structural rearrangement that improves binder accommodation.

Although a similar effect was observed for the 31_RBD binder (Fig. S10, C3), visual inspection of the MD trajectory indicated that this was due to a small rearrangement of its binding interface (data not shown).

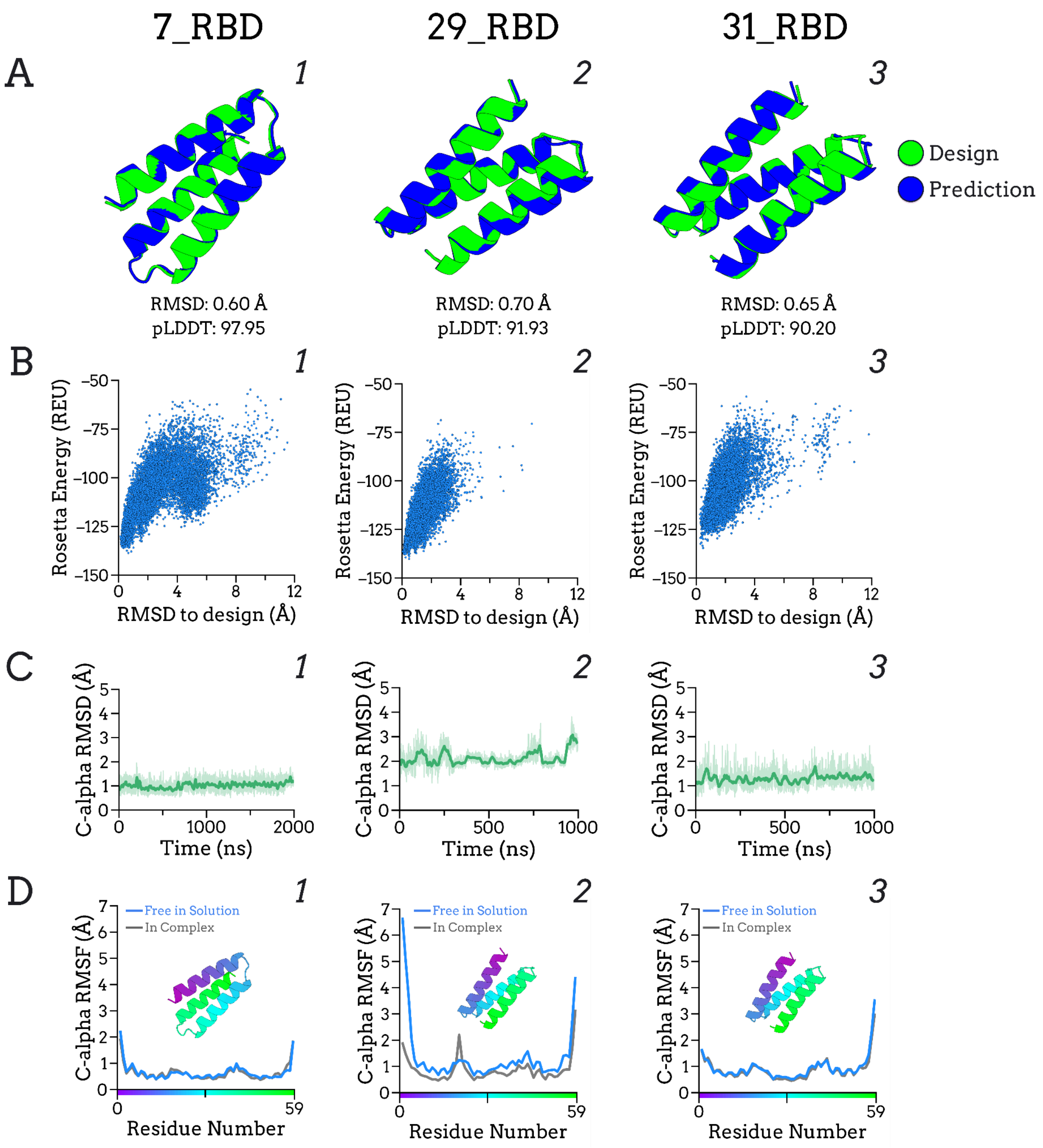


**Figure S11.** Analysis of supplementary SARS-CoV-2 RBD binder monomers. (**A**) AlphaFold2 monomer predictions with designed models (green) superimposed onto predicted structures (blue). (**B**) Folding simulations: Energy versus RMSD plots (**C**) Binder Cα RMSD over time. (**D**) Binder Cα RMSF, both free in solution (blue) and in complex (grey). The 7_RBD system was simulated for 2000 ns, whereas all other systems were simulated for 1000 ns.

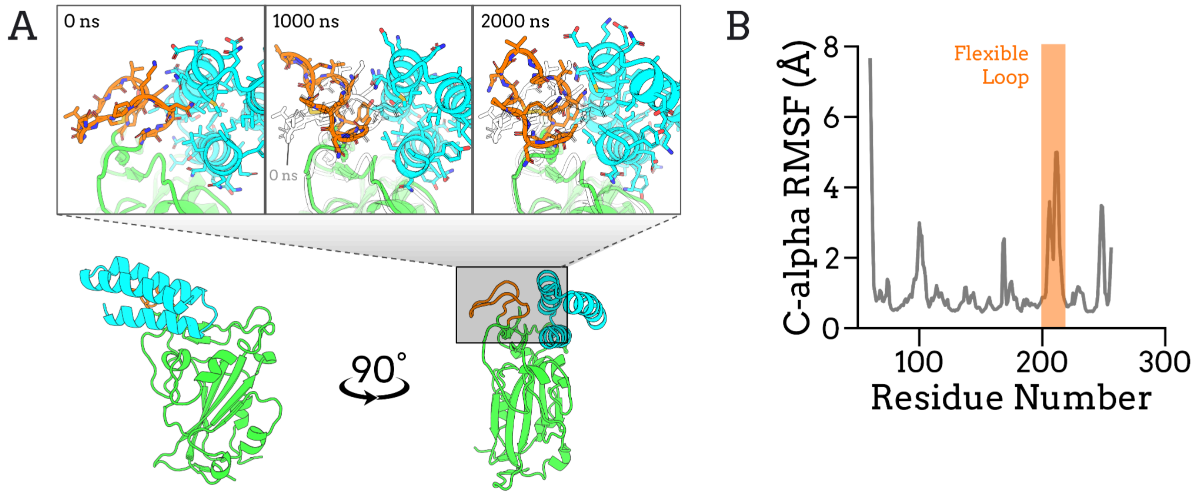


**Figure S12.** Flexible loop dissociation in the 7_RBD complex. (**A**) Simulation snapshots of the 7_RBD complex. The binder is shown in cyan, the RBD in green, and the flexible loop in orange. The initial conformation of the loop is overlaid with transparency. (**B**) Cα RMSF of the RBD during MD simulation in complex with the binder, highlighting the flexible loop in orange, which displays increased flexibility.

#### Section 11: Additional CHIKV and DENV Systems

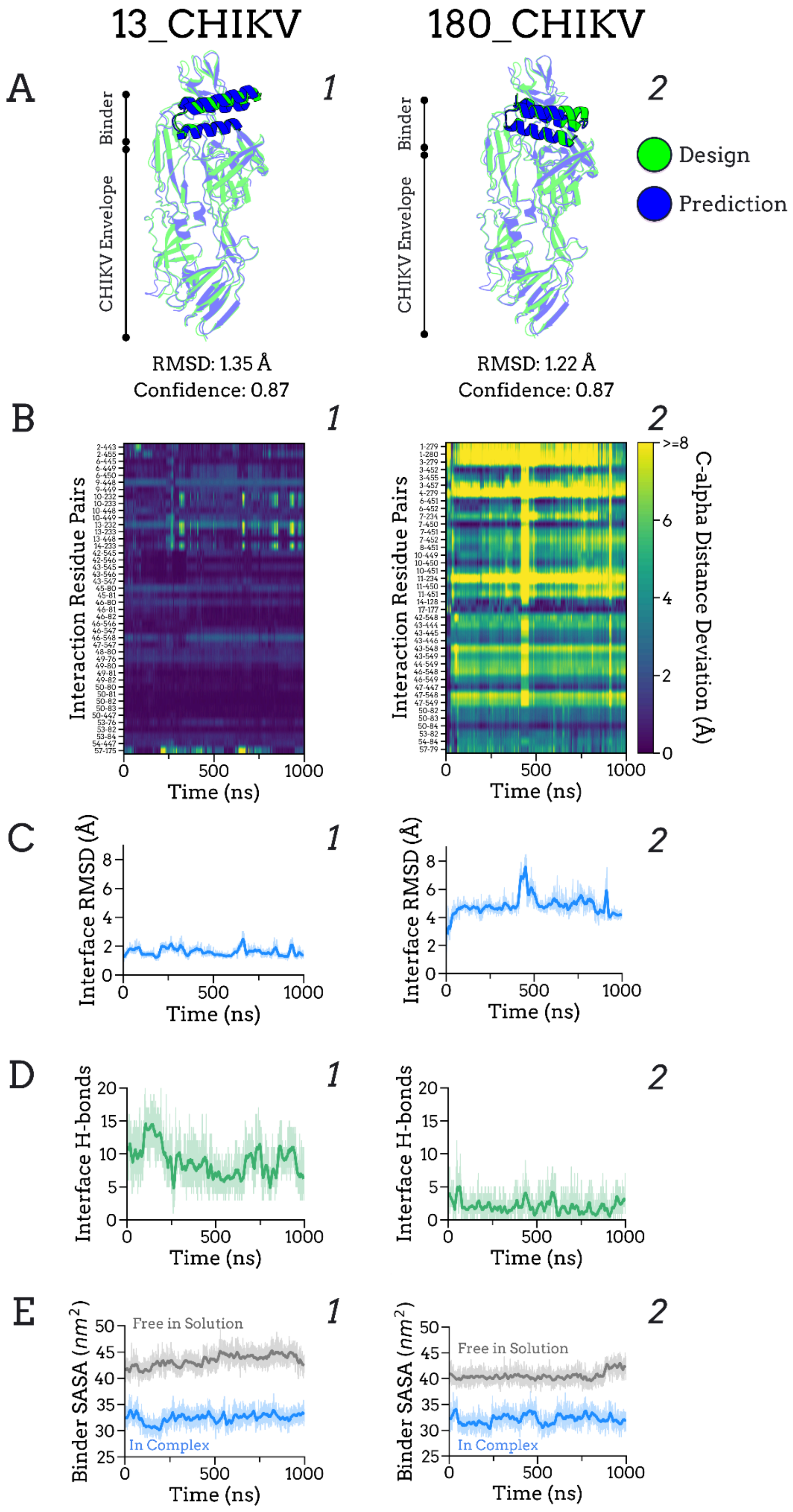

**Figure S13.** Analysis of additional CHIKV binder complexes (**A**) AlphaFold-Multimer predictions with designed models (green) superimposed onto predicted complexes (blue). (**B**) Interaction residue pair stability analysis. (**C**) Interface RMSD over time. (**D**) Number of interface H-bonds over time. (**E**) Solvent Accessible Surface Area (SASA) of the binder in complex (blue) and free in solution (grey) over time.

Interaction residue pair analysis (Fig. S13B) and interface RMSD (Fig. S13C) are reference-dependent metrics (depends on the starting frame), while interface H-bonds (Fig. S13D) and binder SASA (Fig. S13E) are reference-independent. Together they can provide different views of the interaction. For 180_CHIKV, reference-dependent metrics are changing while reference-independent maintain reasonable values, suggesting that although the binder deviated significantly from the designed binding site during MD simulations, it maintained interaction with the target through an alternative conformation. This outcome is not desirable, as the designed binder is expected to interact with the target in the intended conformation.

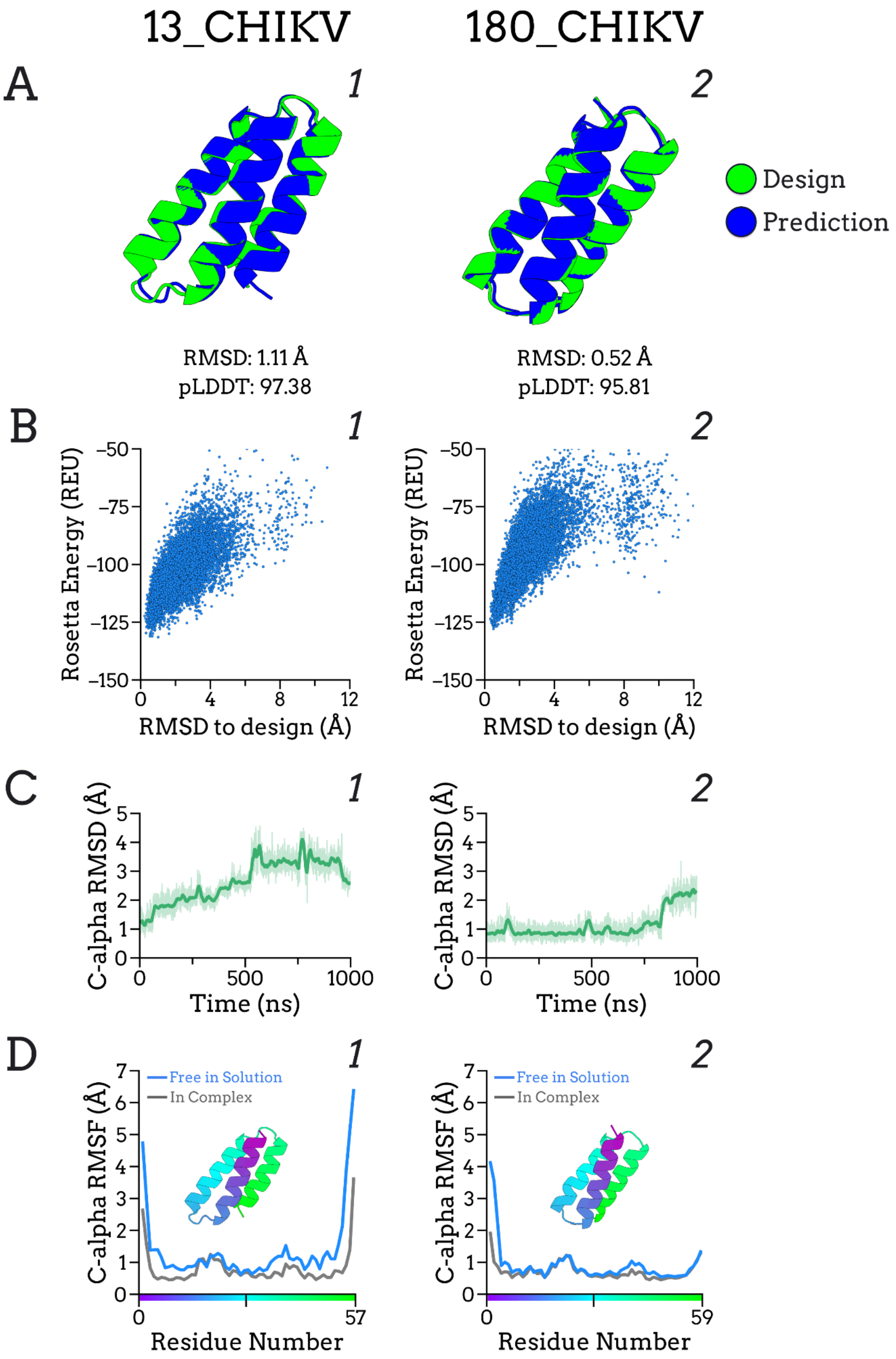


**Figure S14.** Analysis of additional CHIKV binder monomers. (**A**) AlphaFold2 monomer predictions with designed models (green) superimposed onto predicted structures (blue). (**B**) Folding simulations: Energy versus RMSD plots (**C**) Binder Cα RMSD over time. (**D**) Binder Cα RMSF, both free in solution (blue) and in complex (grey).

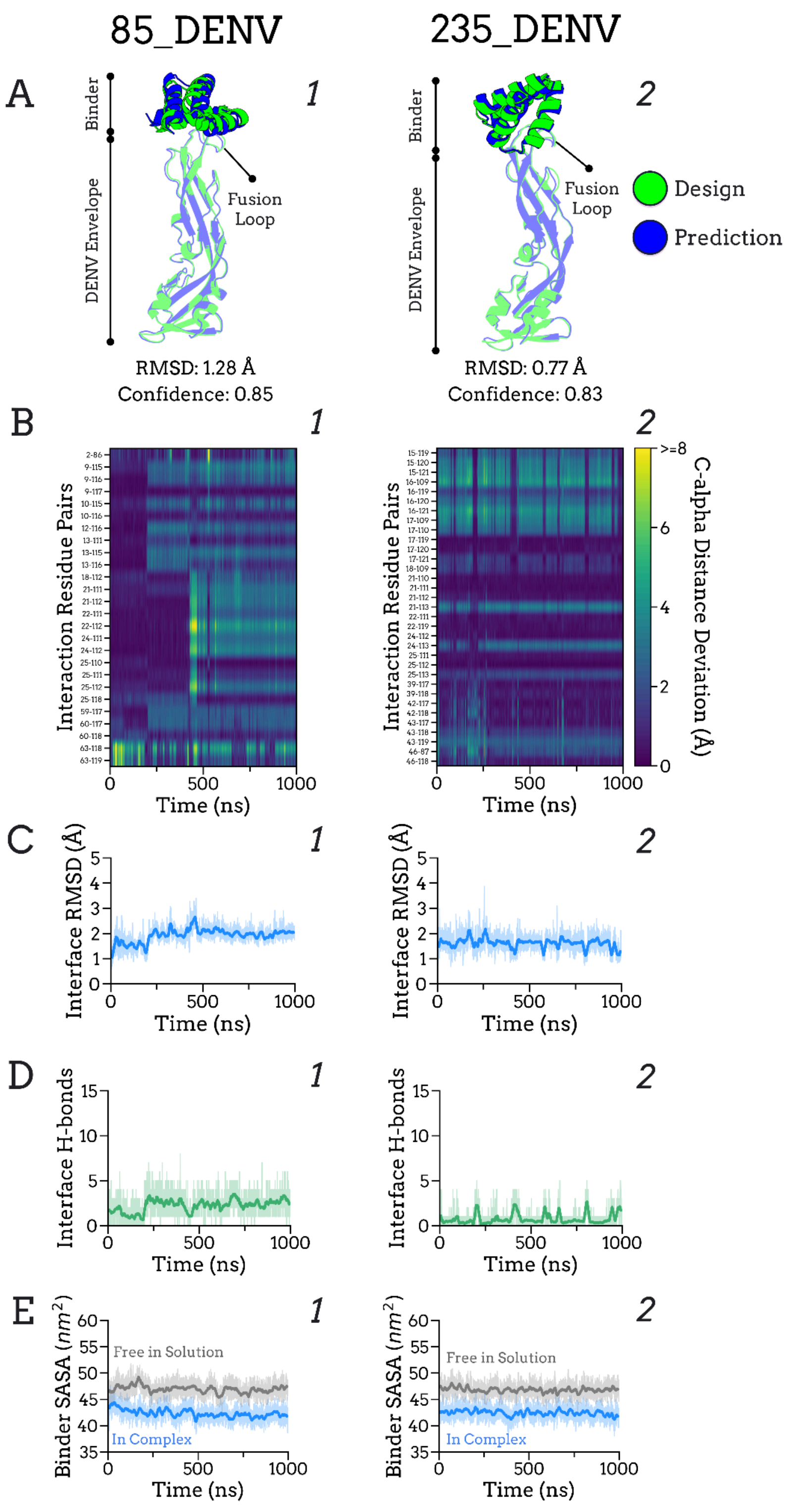


**Figure S15.** Analysis of additional DENV binder complexes (**A**) AlphaFold-Multimer predictions with designed models (green) superimposed onto predicted complexes (blue). (**B**) Interaction residue pair stability analysis. (**C**) Interface RMSD over time. (**D**) Number of interface H-bonds over time. (**E**) Solvent Accessible Surface Area (SASA) of the binder in complex (blue) and free in solution (grey) over time.

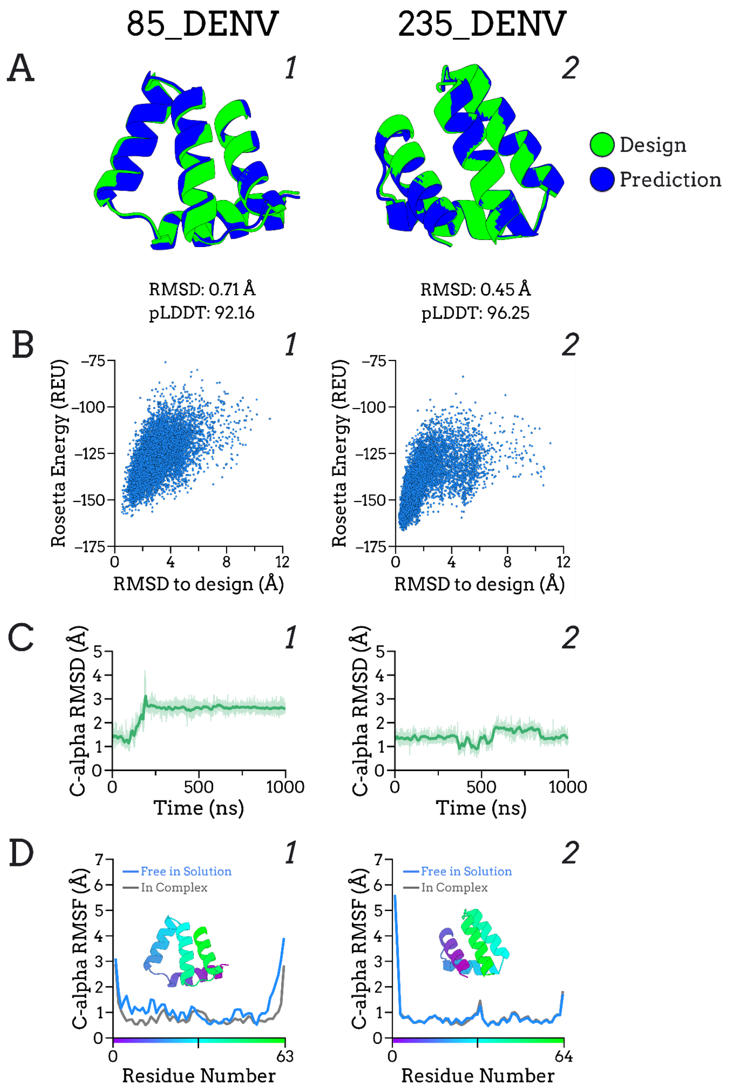


**Figure S16.** Analysis of additional DENV binder monomers. (**A**) AlphaFold2 monomer predictions with designed models (green) superimposed onto predicted structures (blue). (**B**) Folding simulations: Energy versus RMSD plots (**C**) Binder Cα RMSD over time. (**D**) Binder Cα RMSF, both free in solution (blue) and in complex (grey).

A substantial increase in the RMSD of binder 85_DENV during monomeric MD simulations was observed (Fig. S16, C1), followed by stabilization at a higher value, suggesting a possible transition to a more stable conformation. Visual inspection revealed a collapse of the binding residues, effectively closing the interaction pocket (Fig. S17), which would likely impair binding to the target. Although the initial open-pocket conformation may still be accessible through dynamic equilibrium, further analyses would be required to confirm this. As noted in Methods 2.7, such conformational transitions could negatively impact binding.

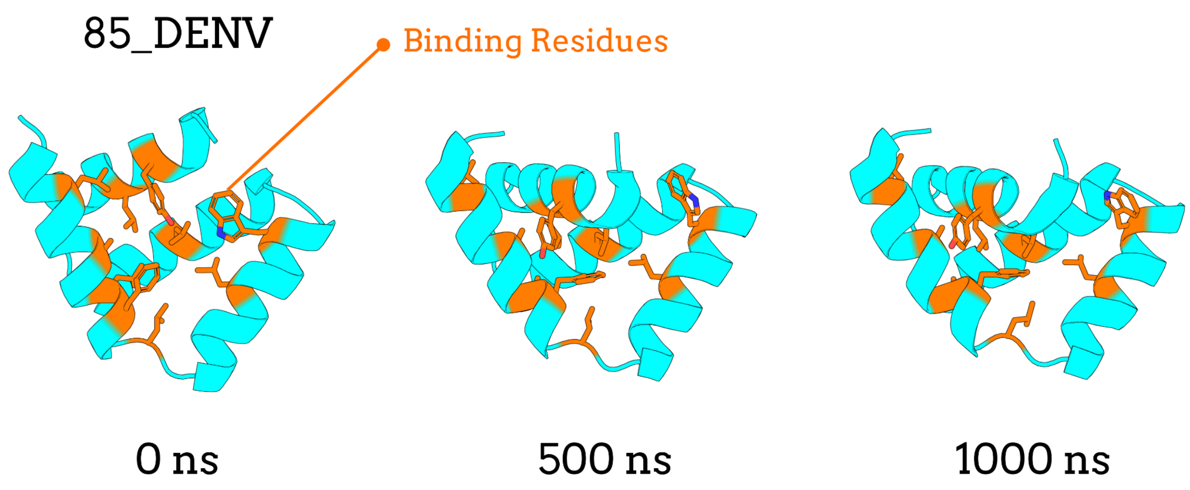


**Figure S17.** Collapse of binding residues in the 85_DENV binder during MD simulations. Snapshots at 0 ns, 500 ns, and 1000 ns are shown, with the binder in cyan and binding residues in orange. Over time, the binding residues undergo a collapse that closes the interaction pocket, potentially impairing target recognition.

| **Supplementary Table 2. Amino acid sequences of analysed binders** | |
|---|---|
| **Name** | **Sequence** |
| 13_CHIKV | SLAAAERALALAVELRNLTDPEEIEARRAEIEALLAQIDDPELRRNVAGQARYIADS |
| **73_CHIKV*** | SRAAIREAEDLYVEAMREGRYLEASHIAALIGRAVQTDDPAEQDALAAEARAHYEAG |
| 180_CHIKV | SDKALADEALQAVNDVLENPTPEVAEPAIRTAEEAIARATNPTVQAIAERALHQVRQLL |
| 85_DENV | SNEVLLAAAEKLFSKLSEEEKQELDELWKADKEAAAEKLRELAKEAGLTEEEREALVEYALKA |
| **98_DENV*** | NVEERVIEKLNEIAKEAPEEKRKELDELFKSDKEAAAEAAAEELLEAGGTEEEAEVAKELVLNA |
| 235_DENV | SEELEKLIEEAAEKLGVSPELLREGLKKLAEDPEALEEYLEKQREAGVDEETLRMIEEAAKLLA |
| **7_RBD*** | SVEEEVENLVEKAKEAIDAGNMEEVNKITNELVNKALNSSDLNDQYLYSKAVNEILNYA |
| 29_RBD | SEAAEVQSAIEALAKAVDAGDKEAVEAEAKKLEELIKKTSDPVVQQVLQEAIEQALKAA |
| 31_RBD | NIKDTLLTLSTNLENAIEAGDKAAVDAVIAEIEKLIAETDDPIIQSNLQAAIDQAKAEI |

**Table S2.** Amino acid sequences of CHIKV, DENV, and SARS-CoV-2 binders analyzed in this study. *Binders shown in bold were those analyzed for an extended time in MD simulations.
